# Collateral mutagenesis funnels multiple sources of DNA damage into a ubiquitous mutational signature

**DOI:** 10.1101/2025.08.28.672844

**Authors:** Natanael Spisak, Marc de Manuel, Molly Przeworski

**Affiliations:** Department of Biological Sciences, Columbia University, New York; Institute of Evolutionary Biology, Barcelona; Department of Systems Biology, Columbia University, New York

## Abstract

Mutations reflect the net effects of myriad types of damage, replication errors, and repair mechanisms, and are therefore expected to differ across cell types with distinct exposures to mutagens, division rates, and cellular programs. Yet when mutations in humans are decomposed into a set of “signatures”, one single base substitution signature, SBS5, is present across cell types and tissues, and predominates in post-mitotic neurons as well as male and female germlines [1–3]. The etiology of SBS5 is unknown. By modeling the processes by which mutations arise, we infer that SBS5 is the footprint of errors in DNA synthesis triggered by distinct types of DNA damage. Supporting this hypothesis, we find that SBS5 rates increase with signatures of endogenous and exogenous DNA damage in cancerous and non-cancerous cells and co-vary with repair rates along the genome, as expected from model predictions. These analyses indicate that SBS5 captures the output of a “funnel”, through which multiple sources of damage result in a similar mutation spectrum. As we further show, SBS5 mutations arise not only from translesion synthesis but also from DNA repair, suggesting that both mechanisms occasionally trigger the same mutagenic process.

---

Mutations fuel evolution, give rise to heritable disease, drive cancers, and contribute to aging [4–6]. In principle, they can stem from many sources. Point mutations and indels, in particular, can occur at any time in the cell cycle, if lesions caused by endogenous or exogenous damage are repaired erroneously. In addition, such mutations can arise during genome replication, from chance misincorporation of a base opposite a canonical nucleotide or from error-prone replication triggered by unrepaired DNA lesions. Identifying which mutagenic mechanism or damage type predominates *in vivo* in any given cell type or tissue is challenging, all the more so in humans.

One approach is to leverage the fact that these sources contribute differentially across cell types and tissues. In cancer genomics, it has become common to model single base substitutions (SBS) in a tumor as a combination of a limited number of “mutational signatures”, then infer the signatures from disparate tumor types. In this representation, each mutational signature is a vector of length 96, with an entry defined as a mutation type (e.g., a base pair transition from C:G to T:A) and the identity of the 5’ and 3’ flanking bases. The idea of the approach, as applied to SBS as well as to double base substitutions and indels [7, 8], is that each signature reflects one or a limited number of damage types, deficiencies in repair, or other processes that operate to varying degrees across tumors [9, 10]. To date, application to data from the COSMIC database [11] has led to the identification of around 100 signatures and helped to link mutation profiles to specific exposures: as one example, signature SBS4 to mutagens in tobacco smoke [12, 13]. While the signatures were originally inferred from mutations in tumors, a subset also accounts for mutations in normal somatic cells and the germline [3, 14, 15].

Two of the single base signatures, SBS1 and SBS5, are found across cancers and in nearly all human tissues and cell types [15–17], including in the germline [3, 14, 15], as well as in other mammals [18, 19]. SBS1 is dominated by transitions at methylated CpG sites, and reflects either spontaneous deaminations or errors in DNA replication [20–23]. Modeling suggests that given these sources, SBS1 mutations should track the number of genome replication cycles [3, 21]. Indirect evidence comes from the analysis of cancers, in which the number of SBS1 mutations increases with age at diagnosis [16] and is higher in metastatic tumors relative to their primary tumor counterparts [24]. In non-cancerous tissues, SBS1 mutations predominate in rapidly-dividing cell types [3, 15], but show little increase with age in post-mitotic neurons, quiescent cells and the female germline [25, 26]. Such observations have led SBS1 to be described as a “mitotic clock” [27, 28].

Like SBS1, signature SBS5 (Figure 1A) is ubiquitous across cell types. Moreover, it is the main signature in a number of somatic cell types, as well as in male and female germlines [ 2, 3, 14, 15, 17, 25]. In contrast to SBS1, the number of SBS5 mutations increases with age not only in dividing cells (e.g., glia) but also in post-mitotic cells (e.g., neurons) (Figure 1B-D), and therefore can arise in the absence of whole genome DNA replication [2, 3]. The etiology of SBS5 remains unknown, with suggestions that it may reflect a collection of endogenous mutagenic processes [15, 17, 29, 30].

**Fig. 1:**
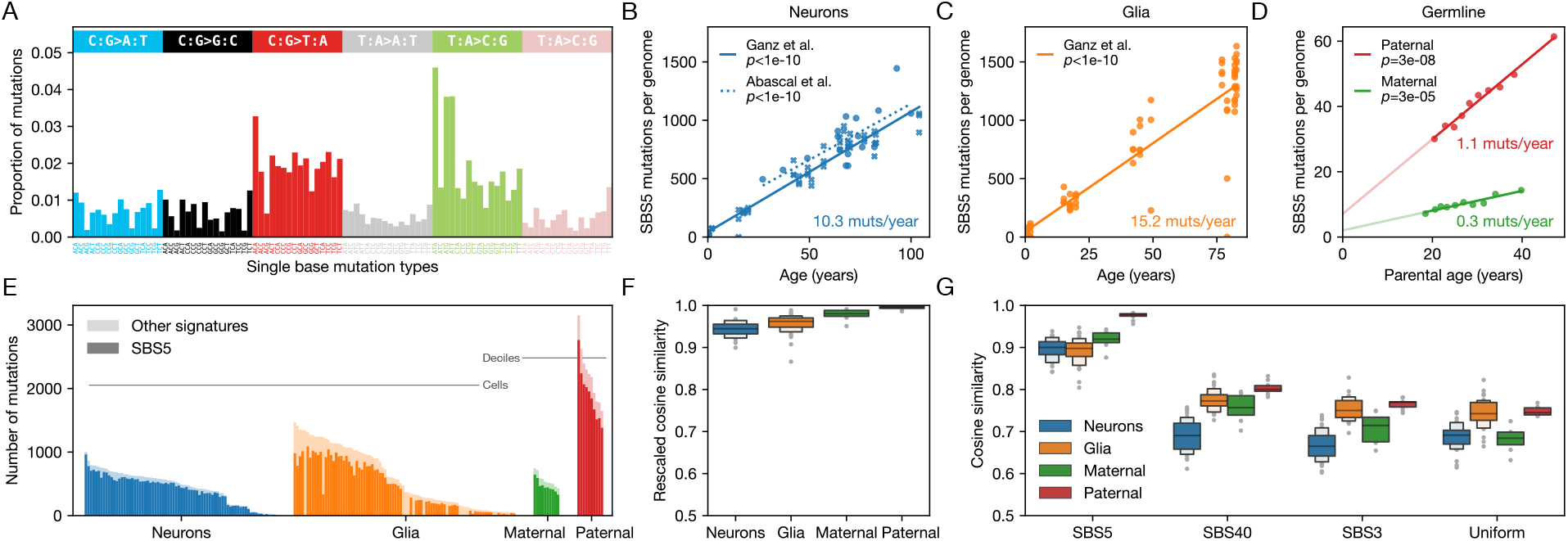
Examples of cell types in which signature SBS5 dominates and provides a good fit to mutation data. (**A**) The distribution of SBS5 mutations over 96 single base mutation types [31]. (**B**) The number of SBS5 mutations increases with age in post-mitotic neurons; data from [25] and [17] are shown with circles and crosses, respectively. Best fit lines and *p*-values are derived from linear mixed-effects models with random intercepts for each individual. Mutation counts per haploid genome (*y*-axis) are adjusted for detection sensitivity [17, 25] (see Methods). (**C**) Same as (B) for glia (data from [25]). (**D**) The number of SBS5 germline mutations increases with deciles of parental age for both maternally-(green) and paternally-phased mutations (red). Data are taken from pedigree sequencing studies and are the subset of mutations phased by transmission in three-generation pedigrees [32, 33]. Best fit lines and *p*-values are from ordinary least squares regression. (**E**) The unadjusted numbers of total mutations (paler shade) and SBS5 mutations (darker shade) detected for individual neuron (blue) and glia (orange) cells, as well as for maternally-(green) and paternally-phased (red) germline mutations (shown in deciles of parental age, as in (D)). Cells and deciles (*x*-axis) are sorted by their total number of mutations (*y*-axis). Data sources are the same as in panels B, C, and D. (**F**) High cosine similarity between observed and reconstructed distributions over 96 substitution types. Shown are the distributions of rescaled cosine similarity. For neurons and glia, we include cells with at least 500 detected mutations. The cosine similarity between the observed and predicted patterns is rescaled by the maximal similarity expected given sampling error (see Methods). Note that the *y*-axis starts at 0.5. (**G**) Cosine similarities after subtracting the contributions of COSMIC signatures other than those we test (i.e., SBS5, SBS40 and SBS3) from the observed distributions over 96 SBS types. Residuals are highly similar to SBS5 and differ more markedly from SBS3, SBS40, or a uniform mutation rate over types. Data sources are the same as in panels B, C, and D. Note that the *y*-axis starts at 0.5.

## SBS5 behaves as a single process

The study of SBS5 presents some technical challenges: for one, its relatively diffuse distribution across possible point mutations (Figure 1A) raises concerns that SBS5 may be hard to distinguish from other “flat” signatures such as SBS3 or SBS40, or that mutations assigned to SBS5 could absorb those that belong to other signatures, and thus that the reconstruction may be distorted by leakage from other processes active in the cell [29, 34, 35]. Reassuringly then, SBS5 provides a good fit to cells in which it is the main signature: for instance, on average, it accounts for 62%-86% of mutations in glia, neurons and paternal and maternal germline (Figure 1E; see Methods); the rescaled cosine similarity between the observed mutations and prediction based on COSMIC signature attribution is 0.94 − 0.99 for these datasets (Figure 1F). After subtracting the inferred contributions of all signatures but those we test (i.e., SBS5, SBS40 and SBS3) from the observed distribution over 96 SBS types, the residual distribution of mutations is substantially more similar to SBS5 than it is to the other flat signatures or to a uniform rate (Figure 1G; see Methods). Moreover, simulations indicate that in data sets of realistic size, the estimated fraction of SBS5 mutations is not sensitive to the presence of SBS16 [27], a signature with which SBS5 often co-occurs (Figure S1). Thus, inferences about SBS5 appear to be robust.

Another challenge in studying SBS5 is interpretative, in that SBS5 could be an amalgamation of distinct endogenous mutational processes that co-occur as a “background” signature (e.g., [29, 35]). Several lines of evidence cast doubt on this possibility, however. First, even across cell types with large numbers of mutations and varying degrees of exposure to mutagens, SBS5 does not decompose into a linear combination of subtypes, in contrast to SBS40 for example [36]. Second, the accumulation of SBS5 mutations with age varies markedly across human cell types [15–17]; as one illustration, it accumulates ~ 0.3 vs ~ 15 extra SBS5 mutations per year per haploid genome in the maternal germline and glia, respectively (Figure 1D and C). For SBS5 to be the superposition of distinct processes would require them to be highly synchronized in their cell-type-specific age effects across all these different cell types. Third, the distribution of SBS5 mutations along the genome is not uniform: for instance, in cortical neurons, SBS5 mutations are enriched in actively transcribed regions, whereas in glial cells, this association is reversed (Figure S2A) [25]. Similarly, SBS5 mutation rates are significantly negatively correlated with large-scale genome accessibility in glia, whereas in cortical neurons, they are enriched in open chromatin peaks (Figure S2B-C). Differences in the distribution of SBS5 mutations have also been reported recently among neuronal types in the cerebellum [37]. If SBS5 were the superposition of distinct processes, such shifts in genomic location would again call for their tight coupling. Therefore, SBS5 behaves as if it were effectively a single process. The question then becomes: what age-dependent process is ubiquitous across diverse cell types?

## Modeling the different modes by which damage leads to mutations

Given that SBS5 is seen across disparate contexts, we reasoned that bringing together observations from tumors, non-cancerous soma and germline under a single framework might help to better understand its etiology. To that end, we developed a mathematical model of mutagenesis that considers the efficiency and accuracy of repair and rates of cell divisions (in the model, we assume that whole genome replication cycles are instantaneous and simultaneous with cell divisions). We build on our previous modeling [3, 21] to explicitly account for collateral mutagenesis and to include two critical features of the interplay of DNA damage and repair: the stochasticity of the damage process and finite repair resources. Concretely, DNA damage is modeled as occurring in stochastic bursts whose magnitude can temporarily exceed the repair capacity of the cell (see Methods for the full model formulation and solutions). In considering the consequences of DNA damage, we distinguish between two mechanisms by which mutations can arise: replication across or near lesions left unrepaired by the time of whole genome replication and errors of DNA repair that occur at any other time point (Figure 2A).

**Fig. 2:**
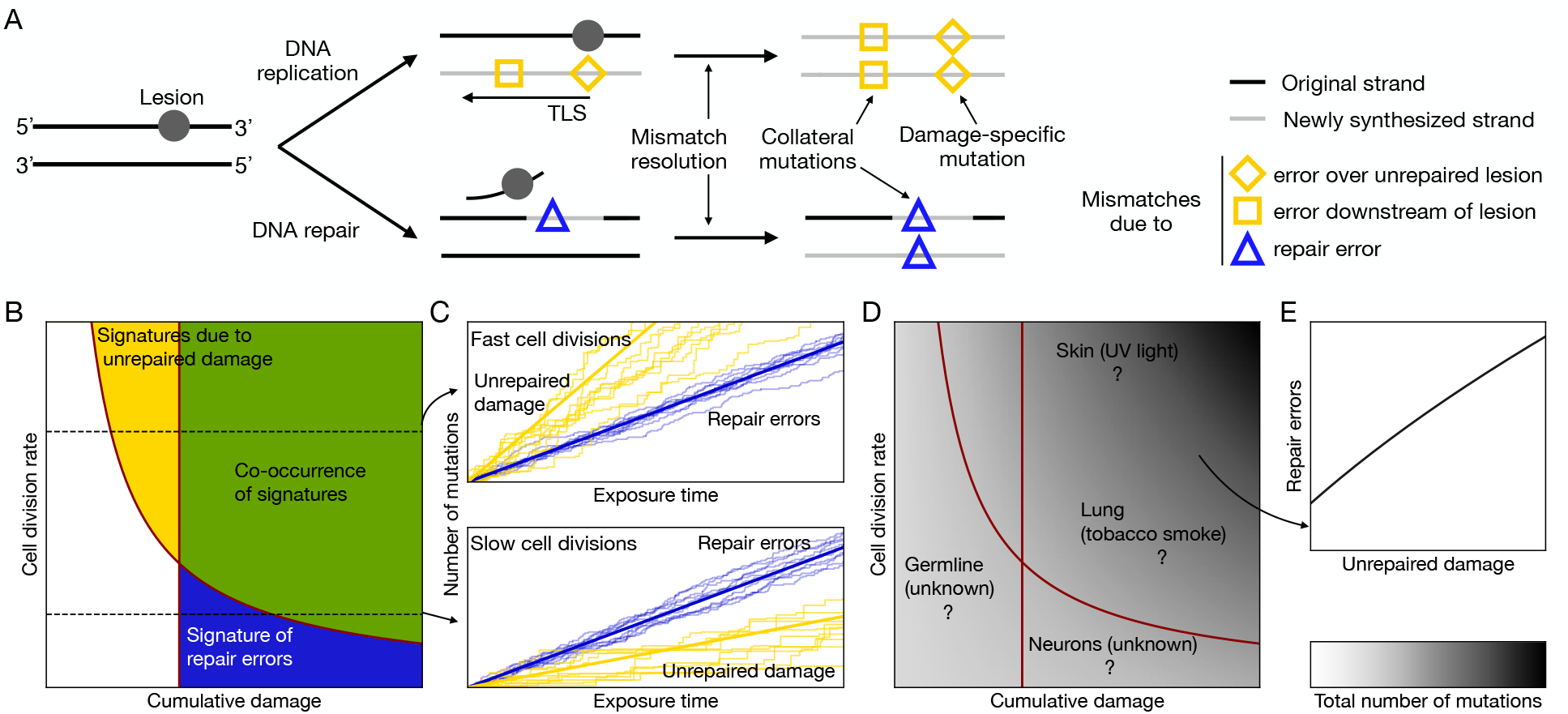
Mutagenesis due to unrepaired and misrepaired damage. (**A**) The cartoon illustrates how a lesion on one of the DNA strands can lead to a mutation at the site of the lesion as well as at a nearby site, thus generating a “collateral mutation”. TLS is triggered by specific lesion types, such as bulky adducts and abasic sites [38]. Repair pathways (pictured), such as nucleotide excision repair, mismatch repair, or homologous recombination, can lead to filling a gap ranging from tens to thousands of bases [39]. (**B**) The type of mutational signatures seen in a cell will depend on the cell division rate and the cumulative damage. Red lines delimit thresholds for signature detection (see Methods). The two types of mutational signatures should co-occur in rapidly dividing lineages with sufficiently high cumulative damage. (**C**) Replication across unrepaired damage is the dominant source of mutation when cell divisions are frequent (top panel) and repair errors when cell divisions are infrequent (bottom panel). The thick lines indicate the expected numbers of mutations due to unrepaired damage (yellow) and repair errors (blue) as a function of the exposure time, which is directly proportional to cumulative damage. Stochastic trajectories simulated from the model are represented by thin lines (see Methods). The blue lines are independent of the cell division rate (see equation 15). (**D**) The total number of mutations per cell expected as a function of cumulative damage and cell division rate. Question marks indicate plausible coordinates of various cell types with known and unknown sources of damage. (**E**) If a source of damage is left unrepaired sufficiently often at whole genome replication and its repair occasionally leads to errors, the numbers of mutations assigned to the two signatures (i.e., the damage-specific signature and the signature of collateral mutagenesis) should be correlated across cells.

Replication across unrepaired DNA damage is error-prone. In some instances, the replicative polymerases can polymerize over the lesion, but the rate of misincorporation is elevated relative to canonical nucleotides: for example, replication over 8-oxo-guanine often leads to a misincorporation of an adenine [40], which contributes to the C:G to A:T substitutions characteristic of oxidative damage and the associated signature SBS18 [41]. For other types of unrepaired DNA damage, such as abasic sites and bulky lesions, the lesion can stall the replicative polymerases and trigger translesion synthesis (TLS), a mechanism of DNA damage tolerance [38]. Because TLS initially recruits low-fidelity inserter polymerases to bypass the lesion, a mismatch can arise directly at the lesion site, represented by a diamond in Figure 2A. For example, TLS over an adduct of benzo[a]pyrene diolepoxide (BPDE) and guanine, a DNA lesion frequently caused by tobacco smoke, leads to the misincorporation of an adenine and contributes to smoking-associated signature SBS4 [13, 42].

DNA damage can also induce “collateral mutations” [43–45], namely mutations that occur near lesion sites as a byproduct of processes of DNA damage tolerance and repair. Such errors can arise when TLS polymerases, after bypassing the lesion, introduce a mismatch downstream (represented by a square in Figure 2A), as extender polymerases synthesize DNA for additional base pairs [46]. Alternatively, collateral mutations may arise through polymerase errors during DNA repair synthesis. For instance, during nucleotide excision repair (NER), a polymerase error during gap-filling can lead to a nucleotide misincorporation either downstream or upstream from the lesion site (represented by a triangle in Figure 2A). These mismatches can then be resolved into a (double-stranded, permanent) mutation during subsequent DNA synthesis, through mismatch repair or during genome replication (the “mismatch resolution” step in Figure 2A). Mutations due to error-prone replication across lesion sites will contribute to mutational signatures that reflect the specific type of damage; in what follows, we refer to such signatures as “damage-specific”. In contrast, because collateral mutagenesis occurs at sites removed from the position of the original lesion, it will lead to a signature primarily shaped by the error spectra of polymerases, rather than by the type of damage.

The model allows us to compare how mutations contributing to damage-specific signatures and signatures of collateral mutagenesis are expected to accrue in different settings. We focus on the regime that is realistic in most *in vivo* contexts, in which the vast majority of lesions are repaired before whole genome replication [47, 48] (see Methods). In this regime, the number of mutations that stem from misincorporations of nucleotides across the lesion or TLS errors downstream from the lesion, denoted *n*_unrepaired_, depends on the balance of the frequency *f* and the mean size of damage bursts *b*, and the rates of repair *r*_1_ and *r*_2_ (which correspond loosely to detection and repair completion, see Methods). This number will accumulate with cell divisions,

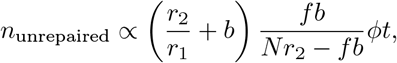

where *ϕ* denotes the cell division rate, *N* the number of repair agents, and *t* denotes exposure time (see Methods). If the cumulative damage, *fbt*, and the cell division rate, *ϕ*, are high enough, there should be detectable signatures of unrepaired damage (yellow and green areas in Figure 2B). In turn, if virtually all lesions are repaired, but the repair machinery makes errors with non-zero probability *ϵ*, the mutations due to such errors will arise at a rate that is independent of cell division rates and will track cumulative damage; assuming a fixed damage rate, their number will be given by *n*_errors_ ∝ *ϵfbt*. If the cumulative damage is sufficiently high, that is *fbt* ≫ *ϵ*^−1^, even non-error-prone polymerases used in repair will lead to a substantial number of mismatches and to a detectable signature of repair errors (blue and green areas in Figure 2B).

A key prediction of the model is therefore that in the regime of high cumulative damage and frequent cell divisions, the same source of damage will give rise to two types of mutational signatures, one due to unrepaired lesions and the other to repair errors (green area in Figure 2B). Across cells, the numbers of mutations contributed by signatures of unrepaired damage should be correlated with one another and with signatures of repair errors (Figure 2E). Moreover, if the same mutagenic process is occasionally triggered by different damage types, then distinct sources of damage will result in a similar signature of collateral mutagenesis. In that sense, and in contrast to the usual interpretation of mutational signatures, collateral mutagenesis will behave as a “funnel”, in which different sources of damage pour into the same mutational output.

## SBS5 tracks multiple sources of DNA damage

We hypothesize that SBS5 is such a funneling signature, explaining its ubiquity across disparate cell types. If so, in cell types exposed to sufficiently high levels of a specific type of DNA damage, the number of SBS5 mutations should increase with the damage-specific signature. With this in mind, we examine the coupling of mutational signatures in whole-genome and whole-exome sequencing of tumors and non-cancerous cells.

In humans, there have been anecdotal reports of associations between the number of SBS5 mutations and damage rates. In lung cancers, the number of SBS5 mutations increases with smoking pack years, a measure of cumulative lifetime tobacco exposure [12]; similarly, in non-cancerous lung epithelia, smokers have a significantly higher SBS5 burden than never smokers [50]. Moreover, in human TK6 cell lines, exposures to temozolomide and several platinum-based drugs significantly increase SBS5 mutation rates [51, 52] (Figure S6). Using the burden of damage-specific signatures as a proxy for cumulative DNA damage, we tested for broader associations in human mutational data. Analyzing mutations attributed to signatures in the Pan-Cancer Analysis of Whole Genomes (PCAWG [53]), we find that correlations between damage-specific signatures and SBS5 are a general phenomenon (Figure 3).

**Fig. 3:**
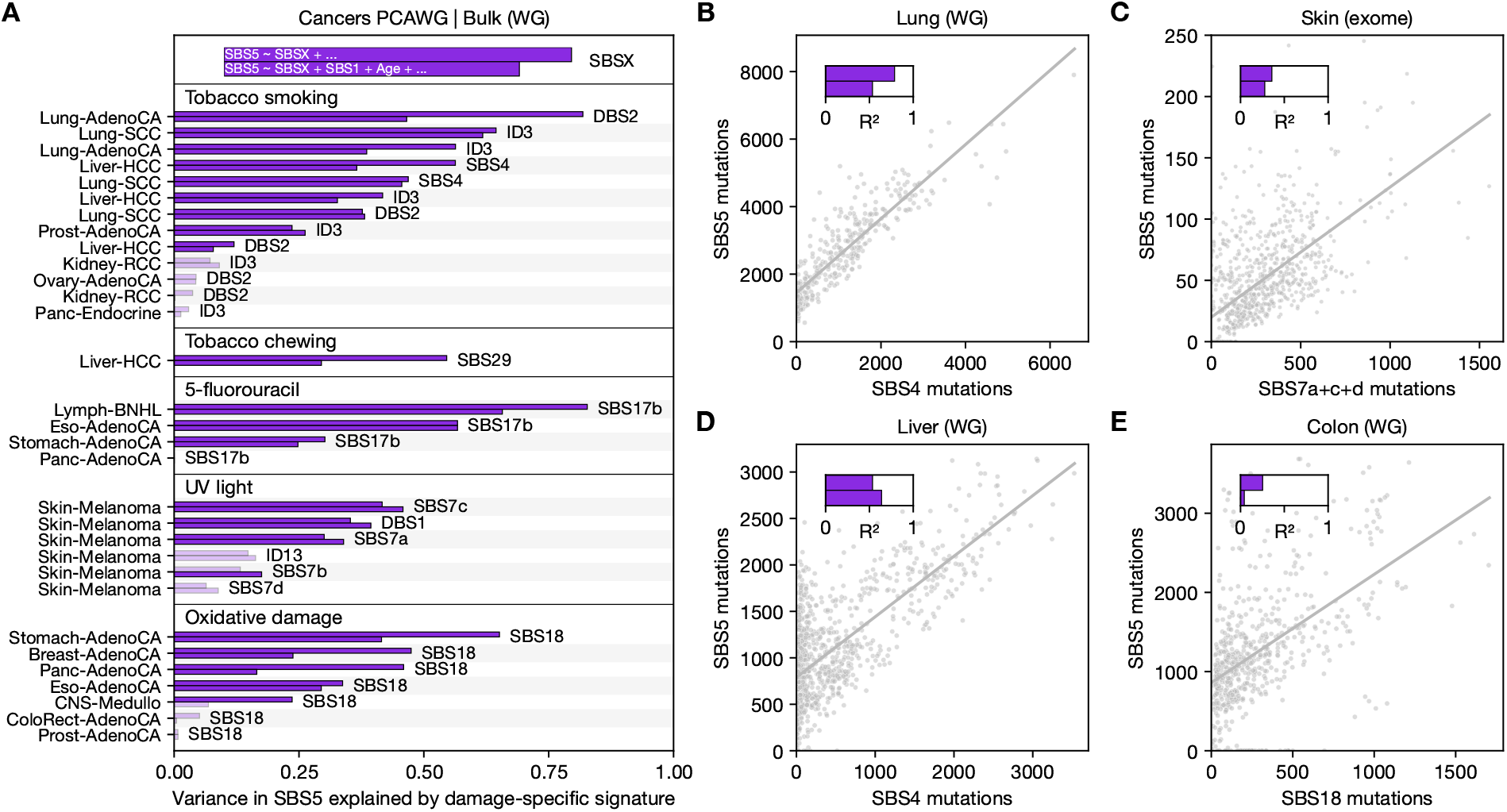
Associations between signature SBS5 and various damage-specific signatures. (**A**) Semipartial *R*^2^ values quantifying the variance in SBS5 attributed to damage-specific signatures across tumors, with cancer types (left), signatures (right), and etiologies (top) indicated. The upper purple bars show the variance in SBS5 explained by a damage-specific signature in a model accounting for two covariates (ploidy and purity; see Methods); lower bars represent the signature’s contribution to variance after also accounting for age and SBS1 (as a proxy for cell division numbers; see main text). Solid bars denote comparisons in which the damage-specific association with SBS5 is statistically significant (i.e., *p <* 0.05 after Bonferroni correction across all tests). The covariates ploidy and purity are excluded from the top legend for readability (but included in the statistical modeling). See Table S1 for data across all tested associations. (**B**) Association between SBS4 and SBS5 in bronchial epithelial cells in smokers. Mutation counts for each signature were attributed using Signet [49], but similar qualitative conclusions are found using alternative methods (Figure S3). The solid line represents the fit to data from all cells. The upper and lower bars show the semipartial *R*^2^ quantifying the variance explained by SBS4 in the models, when regressing SBS5 on SBS4 and SBS5 on SBS4, SBS1, and age, respectively. (**C**) Same as (B), but for liver cells in smokers. (**D**) Association between SBS7a+c+d (sum of SBS7a, SBS7c and SBS7d) and SBS5 in exomes from microbiopsies of skin. SBS7b was excluded due to its bimodal distribution in this dataset, though its inclusion does not change the qualitative conclusion (Figure S4). The solid line represents the fit to the data from all microbiopsies. Purple bars show the semipartial *R*^2^ for SBS7a+c+d, as described in (B). Two outlying microbiopsies, which had more than 250 SBS5 mutations, were excluded from the plot to improve visualization (but are included in the statistical analysis). (**E**) Same as (B), but for SBS18 in epithelial cells of the colon. For a comparison of the observed correlation coefficients reported in (B-E) with simulations that assume all signatures are independent, see Figure S5.

Specifically, we find strong associations between SBS5 and tobacco smoke-related signatures (DBS2, ID3, and SBS4) in lung, liver, and prostate cancers, where these damage-specific signatures account for over half of the SBS5 variance in some cases (Figure 3A). We further detect novel associations between SBS5 and damage-specific signatures linked to tobacco chewing (SBS29 in liver cancer), 5-fluorouracil chemotherapy (SBS17b primarily in adenocarcinomas), UV light exposure (SBS7a/c, and DBS1 in melanomas), and oxidative damage (SBS18 mainly in adenocarcinomas across multiple tissues). In contrast, mutational signatures of repair deficiencies show no or only weak associations with SBS5 (see Figure S7 and Table S1), indicating that correlations among signatures are not introduced spuriously by methods of signature attribution. To confirm these findings, we use an alternative signature decomposition method that was specifically developed to address the challenges posed by flat signatures such as SBS5 [35]. We recapitulate the correlations with all previously mentioned damage sources, with the exception of tobacco chewing, and uncover additional links to damage-specific signatures associated with colibactin (ID18) and the chemotherapeutic agent melphalan (SBS99) (Figure S8). These results establish that SBS5 is frequently associated with damage of distinct types.

The correlations between SBS5 and damage-specific signatures could reflect shared age-related processes and/or numbers of cell divisions leading to the tumors, rather than indicating that the same type of damage led to both types of signatures. Since SBS1 serves as a proxy for cell division numbers [3, 16], we assess whether the associations are substantially weakened when including age and SBS1 mutation counts as covariates in our analysis (lower bars in Figure 3A). For the damage-specific signatures shown in Figure 3, the number of statistically significant associations and the proportion of variance in SBS5 explained remain comparable (Figure 3A). While the associations could reflect other “funneling” processes (e.g., if distinct types of DNA damage cause inflammation that in turn leads to SBS5 mutations), the strengths of the correlations between SBS5 and different damage types in a variety of tumor types support a more direct mechanistic link, as would be expected under collateral mutagenesis (Figure 2E). Also consistent with a causal effect of damage on SBS5, in a mouse model of skin cancer induced by acute exposure to the DNA damaging agent DMBA, the number of SBS5 mutations increases with the DMBA-specific signature [54].

A limitation of bulk sequencing is that it primarily leads to the detection of high-frequency mutations within cell populations. Moreover, while most mutations reported in PCAWG likely predate tumor development [1], it remains uncertain whether associations between damage-specific mutational signatures and SBS5 are also common in non-cancerous cells. We therefore also analyze available non-cancerous datasets where mutations were identified either in single cells [50] or in small clusters of closely related cells [55–57], focusing on single base substitutions, the mutation type that can be most reliably detected in the short-read sequencing data used in these studies. Simulations suggest that with such data, we are able to reliably infer the proportion of SBS5 mutations (see Figure S9).

In people with a history of smoking, we observe a strong association between SBS4 and SBS5 in cells from both the lung (Figure 3B) and the liver (Figure 3C), which persists after controlling for SBS1 mutation counts and age (lower bars in Figure 3B-D). The conclusions remain qualitatively unchanged if we use alternative methods to infer signature contributions (Figure S3; see Methods). Such strong correlations (approximately 78% of the SBS5 variance in lung and 53% in liver cells) are not seen by chance in simulations (see Figure S5). In principle, inter-individual differences (other than a linear effect of age) could contribute to the observed associations in both tumors (Figure 3A) and non-cancerous cells (Figure 3BC). However, in the non-cancerous datasets, where multiple cells are available per individual, linear mixed-effects models incorporating individual as a random effect confirm that SBS4 remains a highly significant predictor of SBS5 counts (*p <* 10^−10^ in lung and liver). Furthermore, phylogenetic relationships among cells within individuals have little to no impact on the associations (Figure S10). Altogether, these findings indicate that smoking-related DNA damage is the dominant factor driving SBS5 accumulation in lung and liver cells of smokers.

To validate the correlation between the UV light-induced signatures and SBS5 observed in melanomas, we analyze exome sequences from skin microbiopsies [57]. Although the relationship appears noisier than for skin melanomas (Figure 3A), possibly due to the much lower mutation counts in exomes, we again uncover an association between cumulative counts of SBS7a+c+d and SBS5 across microbiopsies after accounting for age and SBS1 counts (Figure 3D). A linear mixed effect model with random effects by individual also supports a significant effect of SBS7a+c+d on SBS5 counts (*p <* 10^−10^), and the association persists after removing the effect of phylogenetic relationships among microbiopsies within individuals (Figure S10). Moreover, we infer a significant contribution of UV light-induced signatures DBS1 and SBS7b to the SBS5 burden in whole genome sequences of 40 single-cell derived skin colonies [58], as well as in deep bulk sequencing of 74 genes in skin microbiopsies from 11 individuals [59] (Figure S11). Thus, analyses of mutation data in single cells and small clones corroborate the patterns found in cancer genomes, confirming that the accumulation of SBS5 mutations can arise from distinct damage sources.

Tobacco smoke and UV light primarily lead to bulky lesions, which can trigger collateral mutagenesis due to errors in TLS and/or during their repair (Figure 2A). In colonic epithelial cells, mutation counts of SBS5 are correlated with those of SBS18, a signature associated with non-bulky oxidative damage [56]. We also identify this association in the same dataset, using an alternative signature attribution method (Figure 3E, see Methods). The number of SBS18 mutations explains ~ 25% of the variance in SBS5 across cells (Figure 3E, upper bar in inset), but only ~ 4% after including age and SBS1 as covariates in the model (Figure 3E, lower bar in inset); the decrease is similar in people with and without a diagnosis of inflammatory bowel disease (Figure S12), in small bowel mutations (Figure S12) [60], and across different signature attribution methods (Figure S3). Therefore, in these two tissues, as distinct from a number of tumors (Figure 3A), the correlation of SBS5 with SBS18 appears mostly due to both signatures tracking cell divisions and, to a lesser extent, age (Figure S13 and Table S1). One possibility is that SBS5 and SBS18 are the outcome of related but not identical types of endogenous damage (e.g., bulky versus non-bulky oxidative damage).

In addition to considering associations of SBS5 and damage-specific signatures across cells, we can gain complementary insights by considering their joint distributions along the genome. In smokers, for instance, SBS4 and SBS5 mutation rates are highly correlated across genomic windows; this correlation remains substantial and highly significant after controlling for the first three principal components (which together explain *>* 95% of the variance) of replication timing, genome accessibility, gene density, and GC content (Figure S14A). The same is true for SBS5 mutation rates in non-smokers (Figure S14B). While the correlation between SBS4 and SBS5 could, in principle, arise from two distinct tobacco-induced lesions (e.g., bulky BPDE lesions bypassed by TLS and another type that leads to repair errors), the tight association between both signatures across cells and along the genome suggests a single lesion type.

## The colocalization of mutations with sites of damage points to errors in TLS

SBS5 and SBS4 could arise from a single lesion type in at least two ways: they could both originate during TLS triggered by unrepaired damage or SBS5 could result from repair errors. If both signatures result from TLS, with SBS4 reflecting misincorporations at the site of the lesion [13, 42] and SBS5 secondary errors made by TLS polymerases as they continue synthesis, we should detect clusters of point mutations that occur at nearby positions (diamond and square in Figure 2A, respectively) [45, 61]. We find evidence for such clusters, the vast majority of which are pairs, in the lung and liver of smokers (see Methods): they account for ~ 0.5% of point mutations, of which SBS4 explains 10−12% and SBS5+SBS40 22−64% (Figure 4A; here, SBS5 and SBS40 are combined, because it is challenging to distinguish between them reliably when the number of mutations is very small [49]). The spacing between mutations in a cluster is consistent with the expected length scale of TLS, with an enrichment of pairs separated by *<* 100 bps (Figure 4B) [43, 45].

**Fig. 4:**
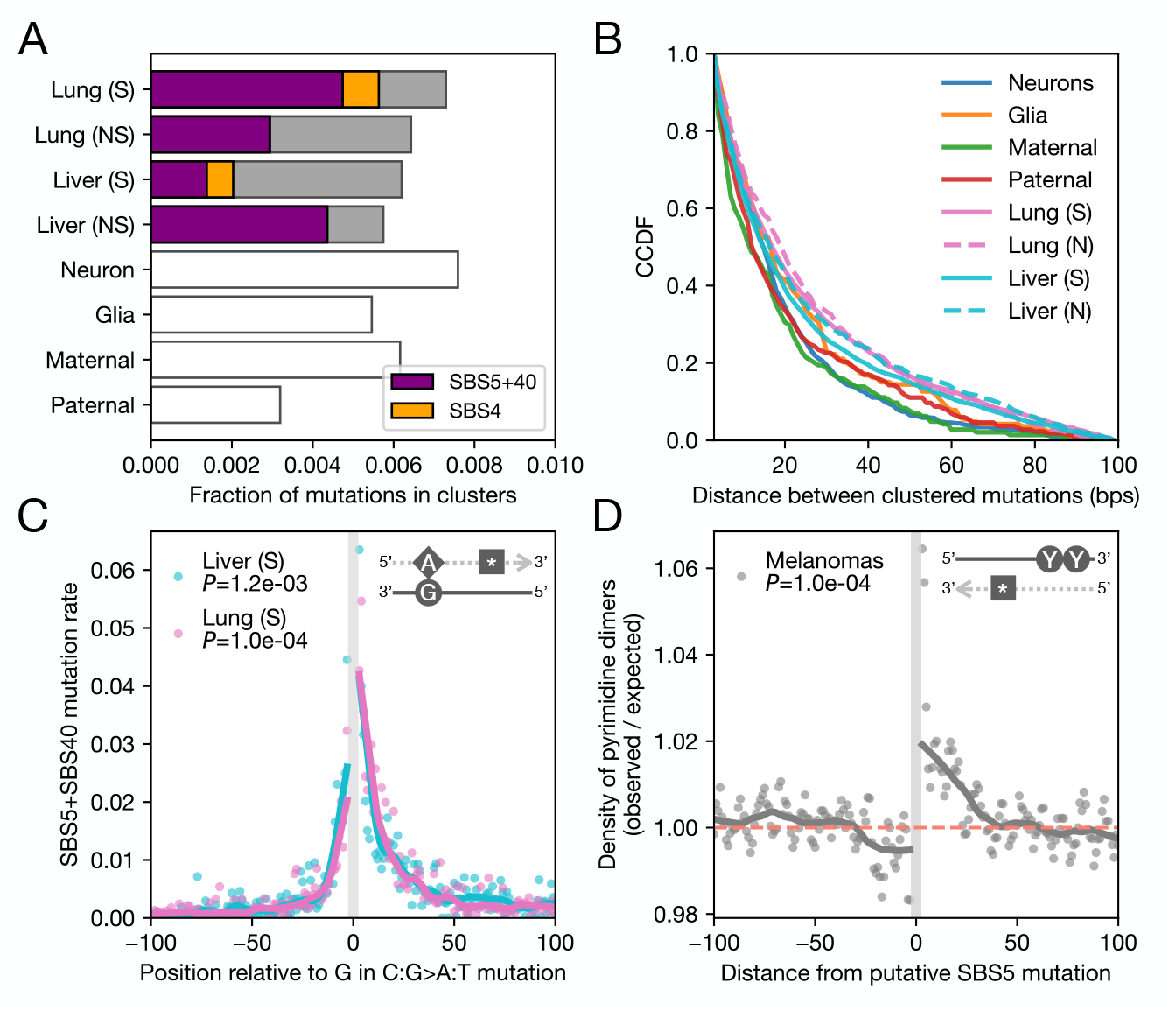
Collateral errors during TLS lead to SBS5 mutations. Throughout, for data from lung and liver, smokers are denoted “S” and non-smokers “N”. (**A**) Fraction of point mutations in clusters across several human cell types. In the lungs and livers of smokers (“S”) and non-smokers (“N”), the proportions of point mutations in clusters attributed to SBS4 (orange) and SBS5+SBS40 (the sum of SBS5 and SBS40; purple) are shown. Due to low mutation counts, signatures were not attributed to clustered mutations in neurons, glia, and germline; the corresponding distributions over 96 SBS types can be found in Figure S15. (**B**) Distribution of distances between mutations within clusters, excluding mutation pairs separated by *<* 3 bps (see Methods). CCDF stands for complementary cumulative distribution function (i.e., 1 − CDF). (**C**) Enrichment in the SBS5+SBS40 mutation rate as a function of distance from C:G to A:T mutation in lungs and livers of smokers. To infer their position, we assume that all C:G to A:T mutations result from a damaged guanine (see main text and inset cartoon, where a diamond represents an error across from the damaged G, the square a secondary TLS error, and the dotted gray arrow shows the newly synthesized strand). Annotated *P*-values were obtained from 10, 000 permutations of distance labels, using the ratio of downstream to upstream density as the test statistic. (**D**) Density of pyrimidine dimers as a function of distance to the nearest putative SBS5 mutation in melanomas. The expected density was obtained by shuffling mutated sites, matching for the original 5-mer context (see Methods). In the inset cartoon, the arrow and square represent the same as in (C), and the circled Ys a damaged pyrimidine dimer. *P*-values were computed as in (C).

Within clusters in lung and liver of smokers, a fraction of mutations are attributed to SBS4 (Figure 4A), within which C:G to A:T substitutions are believed to result from tobacco-induced guanine adducts [13]. We can therefore polarize the clustered mutations with respect to such C:G to A:T mutations and examine where the proximal, putative SBS5 mutation occurs (noting that this polarization will not be perfect, as a subset of SBS5 mutations will also be C:G to A:T substitutions, Figure 1A). As expected if SBS5 mutations in clusters with C:G to A:T mutations arose from TLS, SBS5+SBS40 mutation rates are significantly higher downstream of the putative lesion, in agreement with the direction of DNA replication (*p <* 0.005 in lung and liver; Figure 4C; the two signatures are again combined due to the low number of mutations in clusters). Similarly, we find that pyrimidine dimers (sites typically damaged by UV radiation) are enriched upstream from putative SBS5 mutations in melanomas (*p <* 10^−4^; Figure 4D), in agreement with the associations between UV-induced mutational signatures and SBS5 (Figure 3AC). Together, these findings point to a direct link between SBS5+SBS40 and TLS, consistent with observations of a decreased number of SBS5 mutations in breast cancer cell lines lacking the TLS polymerase REV1 [62] (see also [51]) and a positive correlation between TLS polymerase theta (Pol*θ*) expression levels and SBS5 mutation burden across human cancers [49].

## Evidence for a contribution to SBS5 from errors in repair

A number of lines of evidence suggest that TLS cannot account for all clusters, however. For one, there are comparable proportions of clustered mutations and similar distance distributions in smokers and non-smokers as well as in cell lineages with low rates of divisions, including post-mitotic neurons, and maternal and paternal germlines (Figure 4A-B). These observations establish that mutation clusters can also arise from processes other than TLS, operating at similar length scales (but see Figure S16). The ordering of distance distributions among cell types roughly follows their cell division rates, possibly reflecting different degrees of reliance on TLS (Figure S16).

Moreover, only a minute proportion of all mutations occur in clusters (Figure 4A). For TLS to explain the standalone SBS5 mutations would require TLS polymerases to often bypass lesions without making an error, yet lead to a mutation downstream. Nor can collateral mutations during TLS explain the linear accumulation of SBS5 mutations with age in post-mitotic cell types (Figure 1B).

An alternative possibility, which is not mutually exclusive, is that the widespread association between SBS5 and damage-specific signatures arises from collateral mutations owing to repair errors. If we assume that damage and repair rates are independent along the genome and that the vast majority of lesions are repaired (see Methods), our mutational model makes opposing predictions about these two possibilities. If TLS is the dominant source, we expect SBS5 mutation rates to correlate inversely with repair rates (see equation 18 in Methods). If repair errors are instead the primary cause, the number of SBS5 mutations should correlate with rates of damage but not of repair (equation 19 in Methods).

In the lungs and livers of smokers, DNA damage often arises from bulky BPDE-induced lesions, which are primarily repaired by NER [65]. To test our model predictions, we estimated NER rates, *r*_*m*_, in 5 Mb windows along the genome by fitting the rate of exponential decay of UV-induced lesions over multiple time points, using available data from experiments in a human cell line [66] (see Methods and Table S6).

We then fit to the data the relationship expected from our model, 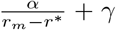 (see equation 20 in Methods), where *α* can be viewed as the sensitivity of mutation rates to repair rates, *γ* corresponds to the mutation rate in regions of high repair, and *r*^∗^ is the critical NER rate below which damage would overwhelm repair. We tested whether *α >* 0, as expected from TLS, and whether *γ >* 0, as expected if there is also a non-negligible contribution of repair errors.

The relationship of SBS4 mutations to repair rates is consistent with mutations arising from error-prone TLS. Estimates of *α* are substantially and significantly above 0 in both the lung and the liver (Figure 5D), whereas *γ* is not significantly different from 0 (Figure 5E), suggesting almost all lesions that could lead to SBS4 are repaired in regions of high repair. For mutation clusters specifically, both *α >* 0 and *γ >* 0 (Figure S17), indicating that a fraction of clusters are caused by TLS, but a subset arises from a process whose activity is correlated with NER along the genome, consistent with our observations for Figure 4AB.

**Fig. 5:**
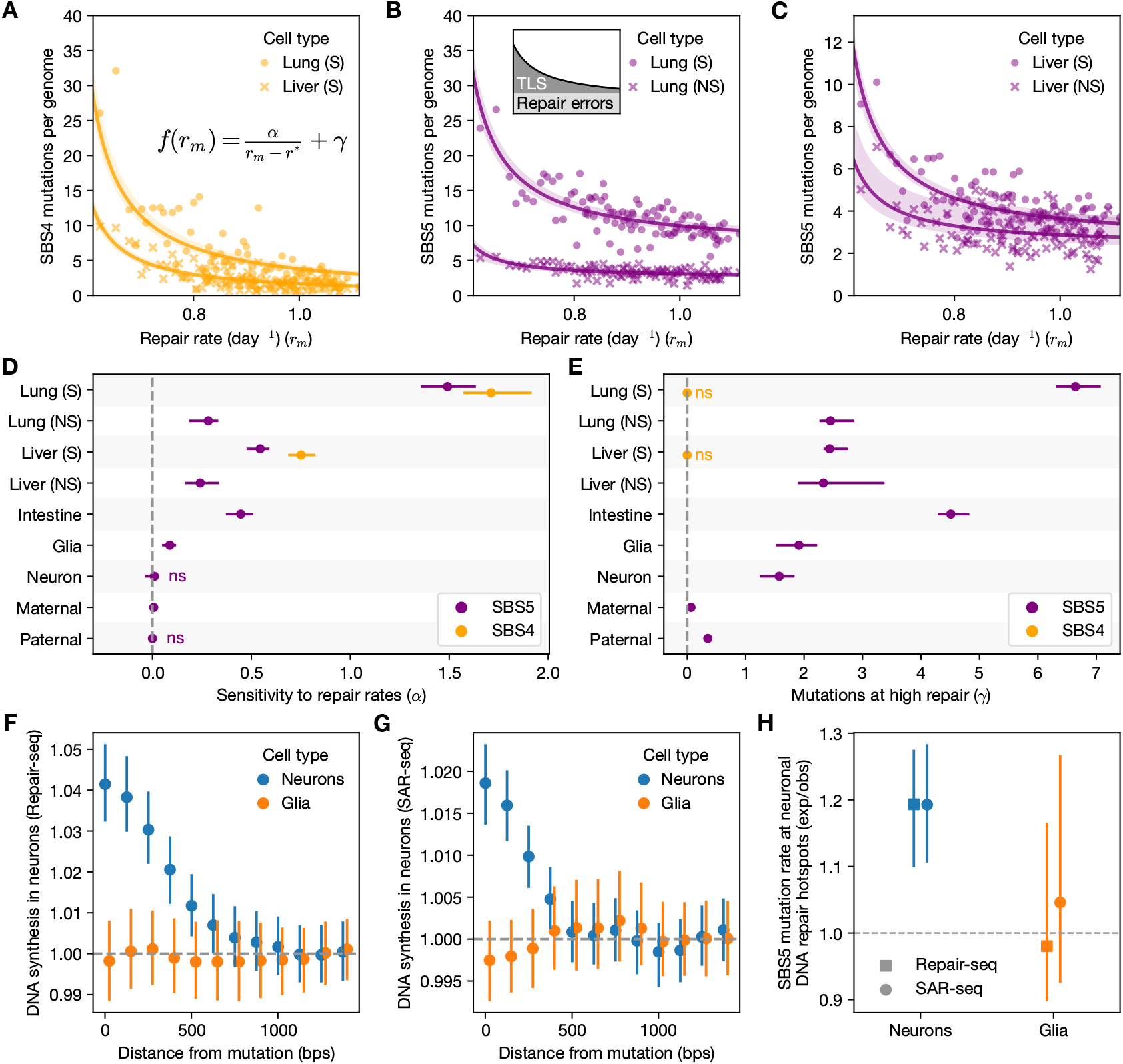
Repair errors can lead to SBS5 mutations. Throughout, for data from lung and liver, smokers are denoted “S” and non-smokers “N”. (**A**) Relationship between rates of NER (*x*-axis, see Methods) and the number of SBS4 mutations per genome (*y*-axis), for lungs (dots) and livers (crosses) of smokers. Each point represents a percentile of the NER repair rate (or 50-quantile for the germline; see Methods), estimated in 5 Mb windows along the genome. Lines represent the fit of the annotated model 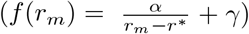. (**B**) Same as in (A), but for SBS5 mutation rates in the lung. Inset: Cartoon illustrating the two sources of SBS5 collateral mutations. The total mutation rate (black line) is decomposed into a constant factor attributed to repair errors (NER) and a TLS component. The colored areas represent the relative contribution of each process across the range of repair rates. (**C**) Same as in (A), but for SBS5 mutation rates in the liver. (**D**) Estimates of *α* across different cell types. 95% CIs were obtained by bootstrapping individuals (or trios) 1,000 times. (**E**) Same as in (D) but for *γ*. We note that the estimates of *α* and *γ* are not independent, as they were jointly fit to the data (see Methods). (**F**) Depth of coverage of Repair-seq [63] as a function of the distance from mutations in neurons (blue) and glia (orange), relative to the average depth at sites *>* 1 kb away from mutations (y-axis). 95% CIs were computed by bootstrapping mutations in each distance bin 100 times. (**G**) Same as in (F) but for reads from SAR-seq [64]. (**H**) SBS5 mutation rates in neuronal DNA repair hotspots based on SAR-seq (circle) or Repair-seq (square) data, for mutations in neurons (blue) and glia (orange). Expected SBS5 mutation rates given the local base composition were obtained by randomly shuffling hotspot positions within the same chromosome and matching for GC content (within 1%) 100 times.

If most SBS5 mutations arise during TLS due to collateral mutagenesis, we would expect to find a similar pattern to that of SBS4. Accordingly, SBS5 mutation rates decrease significantly with repair rates in liver and lung, even after controlling for the principal components of multiple genomic features (Figure S18 and Methods). Rates in smokers are well approximated by 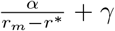 (Figure 5B-C) and estimates of *α* are significantly greater than 0 (Figure 5D). However, in contrast to SBS4, there is a substantial number of SBS5 mutations in regions of high repair (i.e., *γ >* 0; Figure 5E). Furthermore, estimates of *γ* are higher in smokers than in non-smokers (Figure 5B-C), albeit only significantly so in the lung (Figure 5E). These findings indicate that a fraction of the additional SBS5 mutations associated with smoking arise from errors in repair.

In settings in which TLS is rare, there should be little dependence of SBS5 on repair rates along the genome (19). Consistent with this prediction, in neurons, in which the vast majority of SBS5 mutations arose in the post-mitotic state (Figure 1B) [2], the relationship with repair rates is not significant (*α* ~ 0 and *γ >* 0; Figure 5D). The same is true for *α* and *γ* estimates from mutations in the paternal germline; in turn, *α* is only marginally significant for the maternal germline (Figure 5D), possibly due to a relatively large contribution of mutations during early embryogenesis. A modest but significant relationship is found in glia (*α >* 0; Figure 5D), and a stronger one is inferred in the more rapidly dividing colonic epithelial cells, where replication over unrepaired lesions may be more common (Figure 5D). These differences among cell types and tissues are unlikely to arise because *r*_*m*_ captures the repair rates in some cell types better than others (Figure S19) and indeed contrasting *γ* estimates are also obtained for smokers and non-smokers within the same cell type (Figure 5E). Instead, these parameter estimates conform to the predictions of our model, under which the relative contribution of TLS versus repair errors will depend on how often unrepaired lesions persist until genome replication: all else being equal, in cell types with low cumulative damage or infrequent cell division, such as neurons and the germline, SBS5 mutations driven by repair errors are expected to play a more important role (Figure 2C).

To corroborate our findings, we turned to maps of DNA synthesis in post-mitotic neurons, which quantify the occurrence of DNA damage and its repair [63, 64]. Across two independent datasets (Repair-seq and SAR-seq), rates of DNA synthesis during repair are increased near mutations in neurons (Figure 5F-G), of which the vast majority are attributed to SBS5 (Figure 1E). Local base composition effects do not explain these observations (Figure S20; see Methods) and no enrichment of DNA synthesis in neurons is seen near mutations in glia, consistent with previous findings that they occur at distinct genomic locations (Figure S2) [25]. Moreover, when we focus specifically on SBS5 mutation rates in hotspots of repair-induced DNA synthesis in neurons (see Methods) [63, 64], rates in neurons are significantly higher in these regions (~ 20% higher, given their sequence composition; Figure 5H); again, in contrast, mutation rates are not significantly elevated in glia. Together, these results provide further evidence for a role of repair errors in the genesis of SBS5 mutations.

## Discussion

Taken together, our findings indicate that SBS5 is due to collateral mutagenesis originating from two types of mechanisms, TLS during whole genome replication and repair errors arising at any time point. Among the cell types analyzed, we find evidence for both in lung, liver, colonic, and glial cells, whereas in neurons and the germline of both sexes, repair errors appear to predominate. That two types of mechanisms lead to a single signature indicates that at least occasionally, they trigger the same mutagenic process. As even more data accumulate, the slight differences among mechanisms might eventually lead SBS5 to split and reveal the different contexts in which this process occurs.

TLS and repair could both generate SBS5 by a number of possible mechanisms. For instance, when DNA damage leads to single-stranded DNA around lesions (as can happen during TLS and some forms of repair), the exposed conformation may cause the accumulation of further damage [67, 68], the repair of which could result in SBS5 mutations. Another possibility is that TLS and repair can recruit the same polymerase, which funnels disparate lesions into SBS5 mutations after mismatch resolution (Figure 2A). A shared polymerase among pathways could be an evolutionary adaptation to damage tolerance or could arise incidentally, from polymerase kinetics and availability. An error-prone polymerase such as lambda (Pol*λ*) is a possible candidate, given its known role in both TLS and gap-filling of long-patch BER and NER [38, 69]. Polymerase zeta (Pol*ζ*) is also a potential candidate, as it appears to account for a substantial fraction of spontaneous mutations in a human cell line [51] and has been implicated in collateral mutagenesis induced by TLS [70], as well as in the repair of strand breaks [71]. The relatively flat and unspecific spectrum of SBS5 (Figure 1A) and its weak strand asymmetry [31] suggest that a funneling polymerase would have a diffuse error profile, as is known to be the case for an error-prone polymerase (Pol*ϵ*) with a mutated proofreading domain [23].

If repair and tolerance mechanisms recruit a shared polymerase (or trigger another mutagenic process) at different frequencies, a loss of function in a given pathway would either increase or decrease the number of SBS5 mutations, depending on the alternative pathways employed. Differential reliance on this funnel among pathways may help to explain why, e.g., in urothelial tumors carrying missense mutations in *ERCC2*, a key component of NER, the number of SBS5 mutations appears to increase [72] (but see [73]), while in breast cancer cell lines lacking the TLS polymerase REV1, it decreases [62]. Similarly, it may help to explain discrepant observations among mutagen-exposed cells, with *in vitro* experiments leading to increased SBS5 in some cell types [43, 51, 52] but seemingly not in others [42, 74] (Figure S6). In mouse cancer models, there also appears to be a dependence on the mutagen: the number of SBS5 mutations is elevated in tumors induced by DMBA [54] and other carcinogens [75], but not with DEN [76]. Overall, however, the mutagenic process has to be triggered widely enough *in vivo* to generate a ubiquitous mutational signature.

Regardless of the precise mechanism, our finding that SBS5 reflects a funneling step, in which many types of damage, handled by distinct tolerance and repair pathways, result in the same mutational output, explains its ubiquity across cellular contexts and its “clock-like behavior” in many settings [15, 16], i.e., the increase in the number of SBS5 mutations with age. Our results further highlight the difficulty of teasing apart the consequences of exogenous and endogenous sources of damage. An important implication, notably for cancer etiology, is that the burden of mutagens is likely greater than indicated solely by damage-specific signatures.

## Acknowledgments

We thank Ziyue Gao, Scott Keeney, Jonathan Pritchard, Matin Saeidi, Guy Sella, Anastasia Stolyarova, Shamil Sunyaev, and other members of the Przeworski and Sella labs for helpful discussions, as well as Doc Edge, Scott Keeney, William Milligan, Guy Sella, and Anastasia Stolyarova for comments on a draft of the manuscript. We are grateful to Kenichi Yoshida, Matthew Hoare, Tianxiong Yu, Michael Lodato, Joe Luquette, and Christopher Walsh for help accessing data from their publications. This work was funded by NIH grant GM083098 to MP as well as HFSP postdoctoral fellowship LT000257 and Ramón y Cajal fellowship RYC2022-037185-I to MdM.

## Author contributions

NS and MdM performed the computational analyses. MdM and MP supervised the work. NS, MdM, and MP conceptualized the project and wrote the manuscript.

## Methods

### Model of DNA damage

We previously developed a mathematical model of mutagenesis, which includes chance errors made during whole genome replication as well as mutations that arise from damage left unrepaired or repaired incorrectly at any time point [3, 21]. Here, we extend this model in order to focus on two additional and important features of damage and repair: that repair has limited resources and that DNA damage occurs stochastically. By doing so, we are able to study what happens when repair is efficient relative to the typical cumulative damage accrued per cell cycle, as is believed to be the case in most *in vivo* settings [47, 48], but is occasionally overwhelmed by large bursts of damage, and to formulate testable predictions about the behaviors of unrepaired lesions versus repair errors.

In the following, we describe the interplay of damage and repair within a cell and derive how the number of lesions is expected to change over time. We then calculate the expected number of mutations that will result in a lineage of dividing cells. Our notation is summarized in Table 1.

**Table 1:** Parameters and variables of the model.

| Notation | Parameter / variable |
| --- | --- |
| $u(t)$ | Stochastic process of damage |
| $\bar{u}$ | Mean damage rate |
| $x$ | Number of undetected lesions |
| $y$ | Number of repair errors (mismatches) |
| $z$ | Number of free repair agents |
| $N$ | Total number of repair agents |
| $r_1$ | Detection rate |
| $r_2$ | Repair rate |
| $r$ | Effective repair rate |
| $\epsilon$ | Probability of repair error |
| $b$ | Mean size of damage burst |
| $f$ | Frequency of bursts |
| $\phi$ | Cell division rate |
| $p$ | Probability of error-free TLS |
| $\epsilon'$ | Probability of collateral TLS error |
| $q$ | Probability that TLS collateral errors are repaired |

#### Interplay of damage and repair

We consider a source of damage that creates lesions, described by a stochastic process *u*(*t*). To account for the limited resources of the repair machinery, we consider that repair is performed by a finite number of “agents”, denoted *N*, which engage with lesions at rate *r*_1_ (“detection rate”) and disengage when repair is completed, at rate *r*_2_ (“repair rate”). This is a simplified description of multi-step repair pathways such as NER or BER, in which many distinct DNA repair enzymes participate in repair, but captures relevant dynamics for our purposes. The effective parameter *N* can be thought of as the number of enzymes required in the rate-limiting step.

The kinetics of the number of undetected lesions *x* is given by

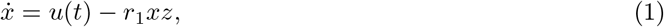

where *z* denotes the number of repair agents available, and we use the dot notation for the time derivative. This number is equal to *N* if no damage is under repair, and decreases when repair agents engage with lesions, *z* ∈ [0, *N*]. Denoting by *r*_2_ the rate at which repair is completed (the expected time for repair to complete after damage is detected is 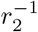), this dynamic can be described as

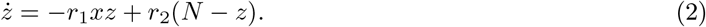

Throughout, we assume that *r*_2_ ≫ *r*_1_ or, equivalently, that we can neglect the few lesions that have been detected but for which repair is not yet completed at the moment of whole genome replication.

We further consider that repair can lead to a mismatch due to a polymerase error. Denoting the probability of such an error by *ϵ*, the dynamics of the number of repair errors *y* are given by

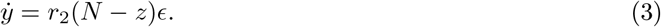

We solve the model at long timescales, i.e. 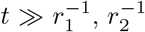, when fluctuations in the number of available repair agents *z* can be neglected, and we assume *z* immediately adapts to the current damage load. In this case, taking *ż* = 0 we find that

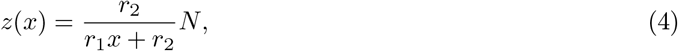

and substituting, we obtain

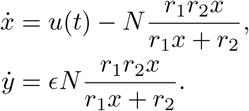

The effective timescale of repair is set by the harmonic mean of the rates at which lesions engage repair agents, *r*_1_*x*, and at which repair is completed, *r*_2_. Introducing the effective repair rate

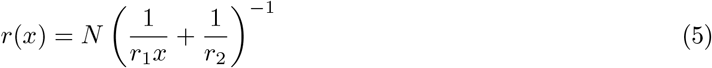

we can now write more simply

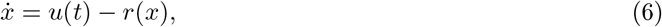

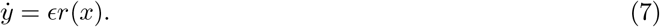

At long timescales, these equations exhibit two distinct dynamical regimes. We characterize them assuming that the damage *u*(*t*) follows a stationary stochastic process with mean 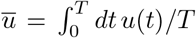, over a sufficiently long window *T*.

First, in the regime of efficient repair, the repair capacity is greater than the mean damage rate over time, i.e., *Nr*_2_ *> ū*. In this case, the number of lesions approaches a steady state that reflects the balance of damage and repair. Alternatively, in the regime of overwhelmed repair, the damage rate surpasses the repair capacity, i.e., *Nr*_2_ *< ū*, and unrepaired lesions accumulate, 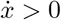.

Next, we calculate the expected number of lesions and repair errors when repair is efficient, and discuss the regime of overwhelmed repair in the subsequent section. The steady-state distribution of the number of unrepaired lesions depends on the distribution of the damage process *u*(*t*). We consider a bursty damage process that can momentarily overwhelm the repair machinery. The bursts of damage are modeled by a Poisson process with rate *f* (the “frequency of bursts”) and the burst sizes are exponentially distributed with mean *b*. To account for the stochasticity in damage, we replace the deterministic equation for the number of lesions *x* (equation 6) with a master equation for the probability density *p*(*x, t*) that the number of lesions is *x* at time *t*,

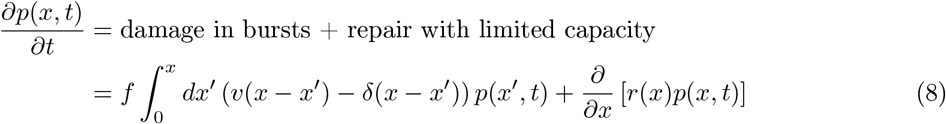

where *v*(*x*) = *e*^−*x/b*^*/b* denotes the distribution of burst sizes and *δ*(*x*) denotes the Dirac delta function. The two terms on the right-hand side describe the probability fluxes associated with damage and repair, respectively. The first term captures damage bursts: probability flows into *x* from states *x*^*′*^ *< x* that are pushed up to *x* by a burst and flows out of *x* as bursts carry it to higher values *x*^*′*^ *> x*. The second term is the deterministic drift due to repair: probability flows into *x* from higher states above *x* and out of *x* toward lower states below, at rate *r*(*x*).

We solve for the steady state *p*(*x*) by setting the time derivative to zero and applying the Laplace transform to the resulting equation, following an approach used for similar master equations that arise in the context of gene expression kinetics [77, 78]. Let *P* (*s*) denote the Laplace transform of *p*(*x*), and *Q*(*s*) that of an auxiliary function *q*(*x*) = *r*(*x*)*p*(*x*)*/x*. The steady-state equation becomes

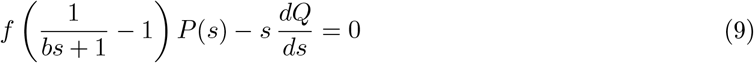

Factoring out *s* and inverting the transform gives an ordinary differential equation of first order for *p*(*x*),

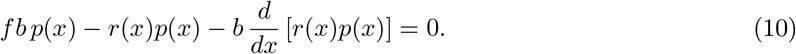

Substituting the explicit form of *r*(*x*) (eq. 5), it can be readily solved to give

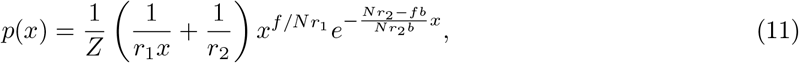

with *Z* a normalization constant. The stationary distribution is therefore given as a weighted mean of two gamma distributions. The expected number of lesions is given by

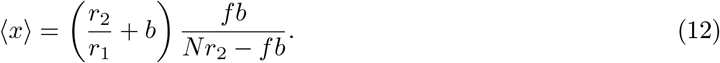

We recover the result of the simpler model from [3], ⟨*x*⟩ = *ū/Nr*_1_ in the limit of *Nr*_2_ ≫ *ū*, i.e., when all damage is repaired before the next burst.

The expected number of repair errors is given by

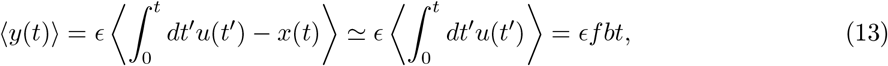

where we assume that at long timescales, i.e., when 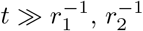, most lesions have been repaired, i.e., 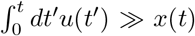, and therefore *ϵ*⟨*x*(*t*)⟩ is negligible. As a result, the number of repair errors *y*(*t*) is a compound Poisson process that tracks cumulative damage.

#### Accumulation of mutations

In this subsection, we calculate the expected numbers of mutations in lineages of cells dividing at rate *ϕ*. To this end, we make the simplifying assumption that whole genome DNA replication is instantaneous and simultaneous with cell division, so that the rate of cell division sets the time to the next whole genome replication. As outlined in the main text, mutations can be caused by unrepaired damage or unresolved repair errors. In the case of unrepaired bulky lesions, the completion of DNA replication requires translesion synthesis (TLS). We denote by *p* the probability that TLS is error-free (i.e., that the TLS polymerase inserts the correct base across from the lesion), by *ϵ*^*′*^ the probability of secondary polymerase error during TLS, and by *q* the probability that such an error is resolved by MMR (this probability may be lower for TLS errors than for the usual errors of replicative polymerases, for which MMR is highly efficient). The average time between cell divisions is *ϕ*^−1^. The number of mutations due to unrepaired damage is then given by

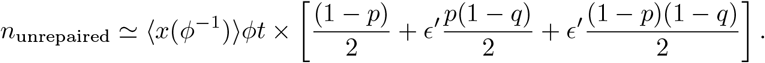

The first term in the bracket is for error-prone TLS, the second for error-free TLS with collateral mutation, and the third for error-prone TLS with collateral mutation (which will lead to a cluster of two mutations). The relative contributions of these three scenarios are determined by TLS characteristics (*p, q, ϵ*^*′*^), and in practice will depend on the type of damage.

To calculate the number of mutations due to repair errors, we assume for simplicity that, overall, half of the mismatches will lead to mutations. This assumption is sensible, given that outside of whole genome DNA replication, MMR is expected to operate symmetrically on both DNA strands [79]. The mismatches that remain unresolved will lead to a mutation in one of the two daughter cells. The number of mutations due to repair errors is thus given by

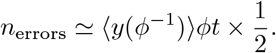

In the regime of efficient repair, the contributions of unrepaired lesions and repair errors to the mutation rate per unit of time are given by

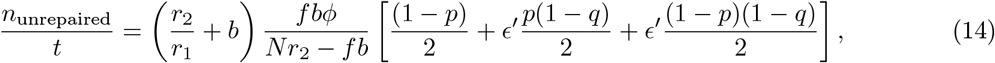

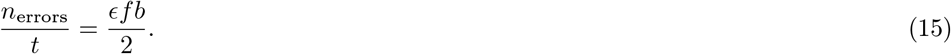

The mutation rate due to unrepaired lesions depends on the balance of damage and repair rates, as well as on the cell division rate. In turn, the mutation rate due to repair errors is independent of repair rates and is simply proportional to the mean damage rate. To draw the phase diagram in Figure 2B, we assumed for simplicity that a fixed number of mutations *n*^∗^ is required to detect a signature (irrespective of its distribution over 96 SBS types or the presence of other signatures), and plotted the detection thresholds *n*_unrepaired_ = *n*^∗^ and *n*_errors_ = *n*^∗^ in red.

#### Dynamics under overwhelmed repair

For completeness, we provide the results of the model in the overwhelmed repair regime, i.e., when *Nr*_2_ *< ū* = *fb*. Unlike in the previous case, the number of unrepaired lesions increases with time,

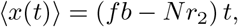

while the rate at which repair errors accumulate is determined by the repair rate, and is independent of the damage rate,

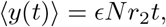

As a consequence, the contribution of repair errors to the total mutation rate will be minimal. If we assume the cell continues to divide despite the accumulation of mutations, the two contributions to the mutation rate will be given by

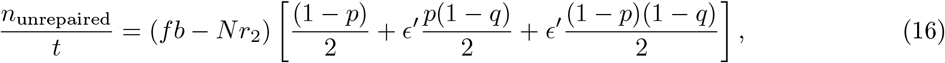

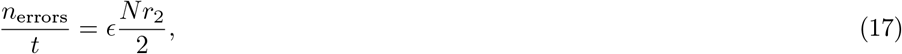

and hence *n*_unrepaired_ ≫ *n*_errors_. *In vivo*, such an uncontrolled accrual of lesions is untenable in healthy tissues and would likely lead to cell death. However, this regime could be relevant for transient or pathological cell lineage behaviors.

#### Simulations

Trajectories of the stochastic dynamics are simulated using the standard Gillespie algorithm, given that the times to the next event (of damage, repair, or cell division) are all exponentially distributed [78, 80]. Briefly, at each step, we draw the waiting time to the next event as an exponential random variable with rate *f* + *r*_1_*xz* + *r*_2_(*N* − *z*) + *ϕ*. The type of the event is then chosen with probability proportional to its relative rate. When a damage burst occurs, we draw the number of new lesions from a geometric distribution with mean *b*. When a repair completes, we draw a Bernoulli variable with parameter *ϵ* to determine whether repair introduced an error. When a cell division occurs, any lesions remaining at that time are resolved according to the replication rules described above.

We used the simulations to confirm the accuracy of our analytic approximation assuming a quasisteady state (4) and to simulate the dynamics described by the original set of kinetic equations (1,2,3); see Figure S21. The code used in the simulations is adapted from [81].

#### Variation in mutation rates along the genome

To interpret the patterns of variation along the genome, quantified in windows *w* of 5 Mb length, we assume for simplicity that all rates (*f, r*_1_, *r*_2_, and trivially *ϕ*) and error probabilities (*ϵ, ϵ*^*′*^, *p*) are the same across the genome. We model the variation of damage rates by varying the expected number of lesions per burst, *b* = *b*(*w*), across the windows *w*. Furthermore, we consider variation in the effective repair rate along the genome, modeled by varying the effective number of repair agents across the windows, *N* = *N* (*w*). For the two modes of mutation accumulation, we then have

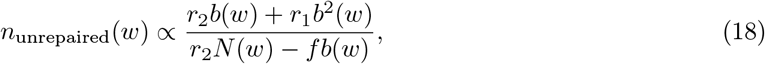

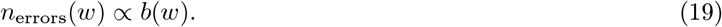

For sources of damage repaired by NER, assuming that damage and repair rates vary independently along the genome, estimates of NER repair rates *r*_*m*_(*w*) relate to the distribution of mutations due to unrepaired damage via

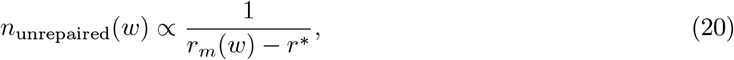

where *r*^∗^ ∝ *fb* is the critical rate, below which the repair is overwhelmed by damage. Importantly, in this setting, the distribution of mutations due to repair errors *n*_errors_(*w*) is expected to be independent of *r*_*m*_.

If, contrary to our assumption, the rates of damage and repair are not independent along the genome, at the scale of 5 Mb, the expected dependence of mutation rates on the estimates of repair, *r*_*m*_, will differ. If damage and repair are positively correlated, the repair errors should be positively correlated with *r*_*m*_ (19); in the opposite case, we expect a negative correlation.

### Data sets analyzed

#### Data from tumor tissues

We obtained mutational signature activities in PCAWG for single base substitutions (SBS), double base substitutions (DBS), and small insertions/deletions (ID), along with clinical metadata (age at diagnosis, tumor ploidy, and purity), from [1]. In addition, we obtained signature attributions in PCAWG using MuSiCal, as provided in [35]. Based on their established etiologies in the COSMIC database (v3.4 [11]), we classified 33 COSMIC signatures as “damage-specific” (Table S2).

### Data from non-cancerous tissues

For neurons and glia, we used mutation data based on sequencing at single-molecule resolution of neurons in [17] and whole-genome amplification of single neurons and glia in [25]. Because these methods do not yield complete genome coverage, we used a global sensitivity of 30% to adjust the mutation burdens in Figure 1B-D, based on estimates in the original publications [17, 25] (to our knowledge, sample-specific sensitivities are not provided).

For maternal and paternal germline mutations, we used whole genome sequencing of trios from three datasets [32, 33, 82]. For all analyses but the inference of clusters, we only considered trios in which over 90% of mutations were phased to maternal or paternal genome. To infer clusters of mutations (data presented in Figure 4CD), we used all phased germline mutations, irrespective of the fraction phased.

For lung and liver tissues, we used whole-genome mutation data from colonies derived from single lung epithelial cells [50] and laser-capture microdissected liver crypts [55]. To investigate the relationship between the tobacco-associated signature SBS4 and SBS5, our analysis was restricted to individuals with a history of smoking.

For skin, we integrated data types from three studies. We analyzed whole-exome mutation data from skin microbiopsies [57], mutational signature attributions from targeted deep sequencing of 74 genes in skin microbiopsies from 11 individuals [59], and whole genome sequences of 40 single-cell derived skin colonies [58]. Because psoralen exposure induces DNA damage that may contribute to SBS5 accumulation independently of UV radiation, we excluded samples with a known psoralen-associated signature from [57].

For intestinal tissues, we used mutation data from laser-capture microdissected colonic crypts [56] and signature attributions from the small bowel [60]. For the colon data, where mutations were mapped to phylogenetic tree branches relating different crypts within an individual, we used the tree topologies provided in the original study to assign mutations back to their respective crypts.

All data sources used are provided in Table S3.

### Attribution of mutational signatures

For PCAWG, we used signature attributions previously inferred using SigProfiler [83] and MuSiCal [35]. For the non-cancer datasets, we relied on attributions provided by the original publications, generated using several tools, including SigProfiler, hierarchical Dirichlet process (HDP [84]), and MutationalPatterns [85] (see Table S3).

In addition, we performed *de novo* assignment of COSMIC v3.3 signatures using SigProfilerAssignment (default settings) on whole-genome mutation data from lung [50], liver [55], and colon [56]. Given that SBS1 and SBS5 could not be confidently separated in exonic skin microbiopsy data using their original approach [57], we inferred signature attribution for this dataset using SigNet, a method better suited for low mutation burdens [49]. We also applied SigNet with default settings to the aforementioned non-cancerous, whole-genome datasets.

To verify whether signature attribution using SigNet is robust and does not lead to spurious associations between signatures, we performed several tests using synthetic data. First, we shuffled the inferred SBS5 and damage-specific mutation counts across samples to remove any underlying correlation; in these synthetic data, SigNet recovers SBS5 counts accurately (Figure S9) and the residual SBS5–damage correlations are significantly weaker than those observed in the real data (Figure S5). We obtained the same conclusion when adding varying numbers of SBS4, SBS18, or SBS7a+c+d mutations to samples with uncorrelated SBS5 counts (Figure S22). Finally, to address possible leakage between SBS5 and SBS16, which share a similar T:A to C:G substitutions pattern, we simulated mixtures of SBS5 with varying proportion of SBS16 (Figure S1) and found that inferred SBS5 loadings are not overestimated.

To quantify the goodness of fit of the signature attribution *P* (*s*), we computed the cosine similarity between the observed distribution over 96 SBS types, *P* (*z*), and one reconstructed from the signatures, ∑_*s*_ *P* (*z*|*s*)*P* (*s*), where *P* (*z*|*s*) denotes the distribution over the 96 types that define a SBS signature. We do not expect a cosine similarity of 1 because of sampling error, which is a function of the number of mutations. In order to account for sampling error, we defined the maximal expected similarity for any given data set as the mean cosine similarity between the observed distribution *P* (*z*) and 1000 bootstrap samples. This quantity provides an upper bound on the goodness of fit using signature attribution. In Figure 1F, we report a rescaled similarity, defined as the ratio of the observed cosine similarity and the expected maximal similarity.

#### Signature attribution along the genome

To attribute mutational signatures along the genome, we aggregated all mutations from a given cell type within a given cohort (e.g., smokers versus non-smokers), and grouped the mutations in 5 Mb genomic windows or percentiles of the repair rate *r*_*m*_ (see section “Estimation of repair rates” below). We attributed signatures using SigNet [49], with an adjustment for mutational opportunities (different 3-mer context content) for each genomic window or percentile of the repair rate. Because the lower number of phased mutations in the germline poses a challenge for reliable signature attribution, we binned the 5 Mb windows for paternal and maternal mutations into 50 quantiles of repair instead of percentiles. To make the estimated values comparable to those reported for other cell types, we divided the number of SBS5 germline mutations in a bin by 2 before fitting the model to percentiles.

### Associations among mutational signatures

To quantify the association between mutations counts attributed to SBS5 and other mutational signatures, we used ordinary least squares (OLS) regression. We quantified the variance explained using the semi-partial coefficient of determination *SR*^2^, which isolates the proportion of total variance in SBS5 counts uniquely explained by a single predictor. This coefficient was calculated as the difference in the model’s *R*^2^ before and after the inclusion of the predictor of interest. For all analyses, predictor and response variables were standardized via *Z*-score normalization prior to model fitting, and statistical significance was assessed using a Bonferroni-corrected *p*-value threshold of 0.05.

In addition, we evaluated partial correlations between SBS4 and SBS5 across 5 Mb genomic windows in lung and liver samples from smokers. To account for potential confounders, we first performed principal component (PC) analysis on four genomic features: replication timing (in H1-hESC cells; UCSC Genome Browser [86]), DNase I hypersensitivity (in GM12878; ENCODE accession ENCFF126OKI), gene density (hg19 annotation; refGene), and GC content (hg19). We then calculated the partial correlations by conditioning on both the first three principal components (PC1–3, which together account for *>* 95% of the variance) and the SBS5 mutation rates observed in non-smokers. This approach allowed us to control for both underlying genomic features and the baseline pattern of SBS5 accumulation expected in the absence of direct smoking-induced damage.

#### Data from tumor samples

For each signature-cancer type combination in the PCAWG cohort, we restricted our analysis to tumors with more than 25 mutations attributed to the non-SBS5 signature, proceeding only if at least 20 tumors remained after this filtering. We fit two OLS models: a baseline model with the non-SBS5 signature counts as the predictor for SBS5 counts, controlling for tumor purity and ploidy as inferred in [87], and a full model that added age at diagnosis and SBS1 mutation counts (as a proxy for cell divisions; see main text) as covariates to the baseline model.

#### Data from non-cancerous tissues and cell types

Analyses were restricted to individuals with multiple samples (i.e., cells or microbiopsies); only samples containing at least one mutation from the relevant damage-specific signature were included. The variance in SBS5 explained by a damage-specific signature was estimated using two models, analogously to the cancer analysis: a model with only the damage-specific signature as a predictor and a second model that included donor age and SBS1 counts as covariates.

To account for the non-independence of samples from the same individual, we tested for statistical significance using linear mixed-effects models. The damage-specific signature and SBS1 counts were included as fixed effects, with a random intercept for each individual. The *p*-value associated with the fixed-effect coefficient for the damage-specific signature was used to assess significance.

To ensure that observed associations were not driven by the genealogical relationships of cells or microbiopsies within an individual, we performed a resampling analysis. We generated 5, 000 replicate datasets by randomly sampling a single cell or microbiopsy from each donor. For each replicate, we recalculated the *SR*^2^ values using the OLS models described above. This procedure allowed us to assess the robustness of the associations in the absence of intra-individual phylogenetic structure (Figure S10).

### Inference of mutation clusters

Point mutations may be in close proximity because they occurred as a result of a single event that led to multiple mutations (such as collateral mutagenesis) or because they arose independently and by chance are near one another. The probability of chance co-localization increases with the number of mutations in a sample and is higher in regions of the genome with elevated mutation rates. Under a uniform mutation rate, in a segment of the genome of length *l*, we expect a triangular distribution of distances *r* ∈ [1, *l* − 1]: out of the 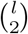 possible pairs of positions, *l* − *r* are separated by a distance *r*, giving

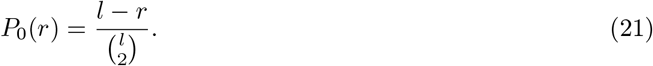

In practice, heterogeneity in the mutation rate will increase the extent of chance co-localization. To characterize what we expect from such heterogeneity, we used the distances between point mutations that occur in independent individuals, and thus cannot have arisen from the same event. We computed the distances within a chromosome arm, to avoid artefactual patterns that arise from lower sensitivity to call mutations around centromeres. Henceforth, *l* denotes the length of a chromosome arm. We excluded mutations at distance *r* = 1, which correspond to double base substitutions (DBS), and *r* = 2, which are often characterized by distinct mutational spectra, possibly due to constraints of sequence context [45], and may be an effect of DBS or consecutive insertions and deletions. In the following, we thus considered *r* = 3, …, *l* − 1.

Across all datasets considered (neurons, glial cells, maternal and paternal germline, lung and liver of smokers and non-smokers), we found relatively good agreement of the distribution of distances between samples with the analytic prediction for a uniform mutation rate (21); see Figure S23A for probability density functions and Figure S23B for cumulative distribution functions for a subsample of chromosome arms (2*q*, 18*q*, 18*p*; chosen to represent different lengths *l*), plotted for all datasets.

Within a sample, we modeled the distribution of distances as a mixture distribution,

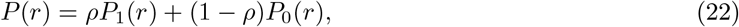

where *P*_0_(*r*) is given by (21), *P*_1_(*r*) stands for the distribution of distances between co-occurring mutations, and *ρ* denotes the cluster prevalence, i.e. the fraction of pairs that are co-occurring. We used the analytical prediction for the null distribution, given that the empirical distribution is noisy at short distances and the analytic approximation provides a good fit (see Figure S23). We assumed a negative binomial form for *P*_1_, which allows us to fit a wide range of possible distributions to the observed distribution of distances. We found the parameters of the negative binomial distribution and the prevalence *ρ*, by fitting a CDF function in a log-log graph (log CDF(*r*) vs log *r*, Figure S24A), using a Levenberg-Marquardt algorithm (scipy function optimize.curve_fit with default parameters). Estimates of prevalence across cell types are shown in Figure S25. In all cell types, there is evidence for co-occurring mutations, i.e., our estimate of *ρ >* 0, and the mixture distribution (22) provides a good fit to the data. From the fit, we estimated the expected false discovery rate for a given threshold distance *r* (Figure S24B), i.e., the fraction of mutation pairs misclassified as co-occurring, when they in fact only co-localize by chance,

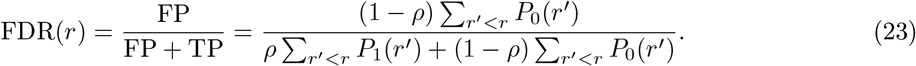

For consistency, we chose the same threshold of *r* = 100 across all datasets, and excluded data from chromosome arms with FDR(*r* = 100) *>* 5% (Figure S24C). Clusters are therefore defined as mutations within 100 bp of one another. Because 98.5% of clusters consist of only two mutations, we restricted the downstream analysis to co-occurring pairs.

We attributed signatures to mutations in clusters found in lung and liver, in smokers and non-smokers separately (clustered mutations were not assigned to signatures in neurons, glia, and germline due to low mutation counts). Given that only ~ 0.5% of mutations fall in clusters (Figure 4A), we pooled mutations from all samples within a cohort and attributed signatures using SigNet, obtaining loadings *P* (*s*). Signature attribution yields a high goodness of fit to the observed 96-type distributions in all four cohorts (cosine similarities to observed mutation profiles ≥ 0.95). SBS5 and SBS40 were combined, because it is difficult to distinguish between them given the small numbers of mutations [49]. To verify whether heterogeneous pairs of mutations, where one is attributed to SBS4 and the other to SBS5/40, are present among the clusters, we estimated a pairwise signature density *P* (*s*_1_, *s*_2_). To do so, we iterated over all pairs of mutations in a given cohort, and summed the probabilities that one of the mutations is due to signature *s*_1_ and the other due to *s*_2_,

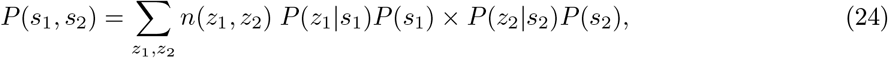

where *z*_1_ and *z*_2_ go over 96 SBS types, *n*(*z*_1_, *z*_2_) denotes the number of pairs in which one mutation is of type *z*_1_, and the other of type *z*_2_; *P* (*z*|*s*) denotes the distribution over the 96 types that define a SBS signature. This pairwise density is reported in Figure S26; it confirms that a substantial proportion of clusters are heterogeneous and correspond to co-occurrence of SBS4 and SBS5/40.

### Patterns of sequence composition around mutations in melanoma

We downloaded mutations in melanoma identified by the PCAWG [87]. Because most mutations in this cancer type are due to UV radiation (SBS7a/b/c/d), in order to focus on putative SBS5 mutations, we restricted our analysis to mutation types that account for *<* 1% of mutations in these COSMIC UV signatures. Among mutations not explained by SBS7a/b/c/d, 72% are attributable to SBS5, while SBS40 and SBS1 account for 18% and 4%, respectively [1].

To account for effects of local base composition, we shuffled the positions of the original mutations within 250 kb of the original location, matching for their 5-mer context. For both observed and shuffled mutations, we quantified the density of pyrimidine dimers (CC, TT, CT, and TC) as a function of distance from the mutated site (up to ±100 bp), folding the density of complementary purine dimers (GG, AA, GA, AG) onto the top strand. To determine whether pyrimidine dimer density was significantly asymmetric around the mutation sites, we computed the ratio of the total dimer density downstream to the total dimer density upstream. Statistical significance was assessed by permuting distance labels 10,000 times.

### Estimation of repair rates along the genome

To estimate NER rates along the genome, *r*_*m*_, we used genome-wide maps of UV-induced cyclobutane pyrimidine dimers (CPDs) across six time points (0, 1, 8, 24, 36, and 48 hours), based on Damage-seq in an experiment in the human cell line GM12878 (GEO Accession: GSE98025; [66]). In this regard, we note that damage rates cannot be estimated using existing genome-wide maps of BPDE lesions taken at a single time point, because repair starts removing lesions within cells shortly after BPDE exposure, such that the signal of damage and repair is conflated [13]. For each 5 Mb genomic window, we estimated the exponential decay of CPD lesions by fitting a linear regression to the log-transformed lesion counts (ln(count + 1)) as a function of time. The estimated repair rate *r* was derived from the negative slope of this fit and scaled to represent the rate of repair per day. For some of the analyses (e.g., Figure 5), we grouped the windows in percentiles of *r*_*m*_.

To compare the association between *r*_*m*_ and genome accessibilities across different cells/tissues (Figure S19), we analyzed DNase I hypersensitivity data generated by ENCODE for testis (ENCFF589CRI), liver (ENCFF606YDZ), lung (ENCFF787QGK), brain (ENCFF331PTE), colon (ENCFF231JLT), and the cell line GM12878 (ENCFF126OKI). We estimated the genome accessibility in each dataset by summing the product of the signal intensity and the region length for all annotated hypersensitive sites within each 5 Mb genomic window.

### Association between repair rates and mutation rates

To test the relationship between repair rates and mutation rates along the genome, we modeled the mutation burden as a function of the local repair rate by fitting (cf. equation 20 above):

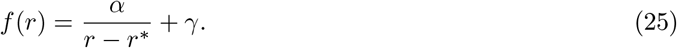

We estimated *α* and *γ* for each mutational signature and cell type using non-linear least squares optimization. We fixed *r*^∗^ = 0.55 day^−1^, as similar estimates were obtained across datasets when this parameter was allowed to vary. To estimate uncertainty in our parameter estimates, we performed 1,000 bootstrap resampling iterations across individuals (or trios) within each cell type. The 95% confidence intervals for *α* and *γ* were then determined from the 2.5th and 97.5th percentiles of these bootstrapped distributions.

### Relationships between mutation rates and DNA synthesis in post-mitotic neurons

To quantify repair-induced DNA synthesis in post-mitotic neurons, we analyzed Repair-seq [63] and SAR-seq [64] data (SRA accessions listed in Table S4). Raw sequencing reads were aligned to the human reference genome (hg19) using BWA-MEM [88]. To assess the levels of DNA synthesis surrounding mutated sites, we quantified depth of coverage across 250 bp sliding windows with a 125 bp step size in regions ±2 kb away from each mutation. We focused on mutations in neurons (from [25] and [89]) and glia (from [25]) (Table S3). To estimate 95% confidence intervals, we performed 500 bootstrap resampling iterations across windows for a given distance. To allow for direct comparisons between neurons and glia, we computed relative DNA synthesis by dividing the read depths by the mean read depth in windows more than 1 kb away from mutations.

To determine whether SBS5 mutation rates differ at DNA repair hotspots based on Repair-seq and SAR-seq peaks, we generated 100 synthetic sets of control regions by randomly shuffling the original peak coordinates within the same chromosome. To control for local base composition effects, we ensured that the GC content of each synthetic region matched its corresponding original peak within a ±0.5% margin. We then inferred signature attributions for mutations in neurons and glia using SigNet across all these sets, adjusting for mutational opportunities as explained in “Signature attribution along the genome”. We computed 95% confidence intervals for the ratio of observed to expected SBS5 mutation rates by taking the 2.5th and 97.5th percentiles across the replicates.

## Data and code availability

The sources of the mutation data are listed in Table S3. The code to reproduce our analyses is available on GitHub at https://github.com/n-t-n-el/sbs5.

## Supplementary information

### Supplementary figures

**Fig. S1:**
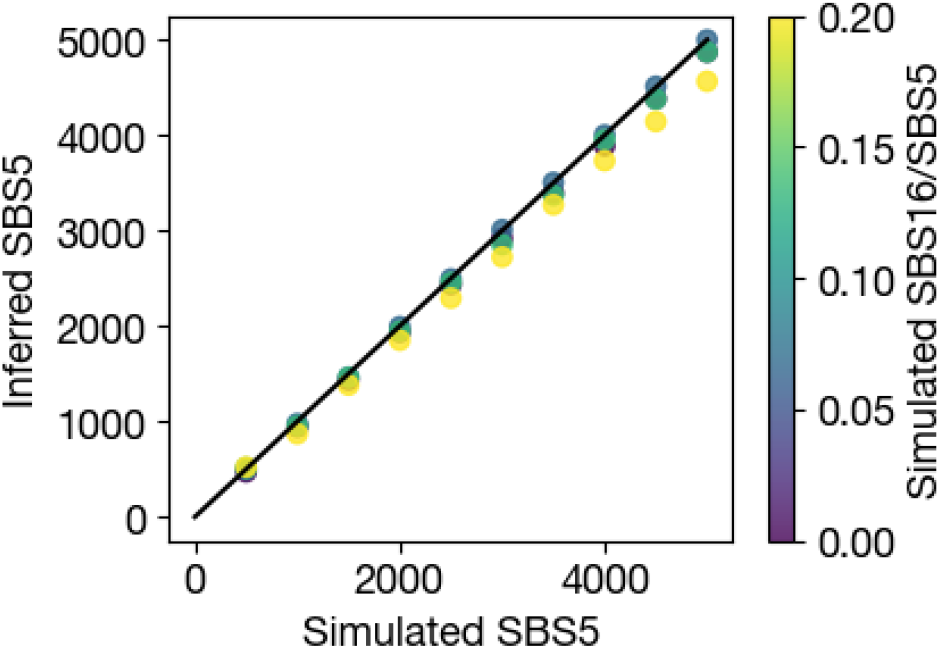
Comparison of simulated and inferred SBS5 mutation counts. In the simulations, SBS5 mutation counts vary between 500 and 5000, matching the mutation rates in the datasets used in our study. The number of additional SBS16 mutations varies as a proportion of SBS5 mutations, between 0 and 20%. For each combination of SBS5 and SBS16 mutation counts, we ran 20 replicates and inferred signature attribution using SigNet. Points show the mean across replicates and the color scale reflects the changing proportion of additional SBS16 mutations. The inference of SBS5 rates is robust to the presence of signature SBS16: the inferred loading of signature SBS5 is never over-estimated and only slightly under-estimated under highest rates of SBS16 (SBS16/SBS5 = 20%).

**Fig. S2:**
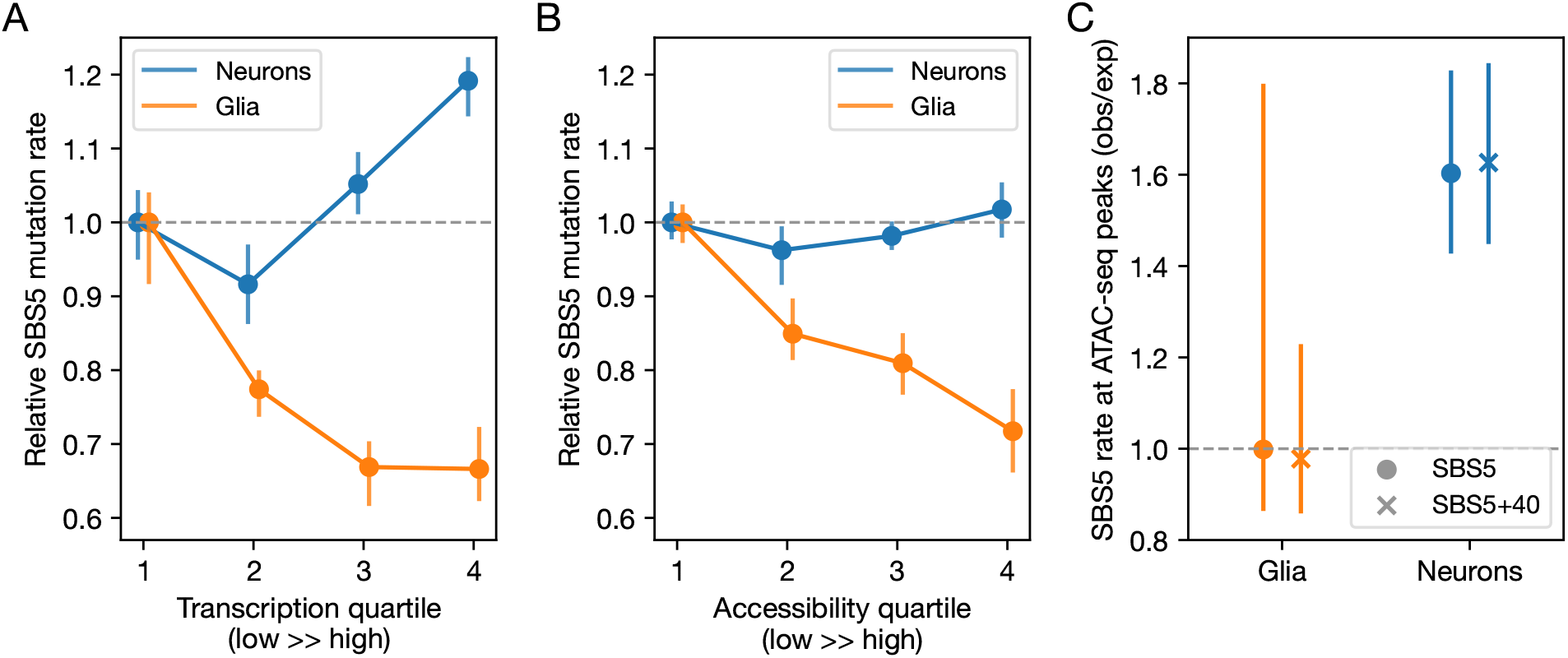
Distinct relationships to SBS5 mutation rates in neurons versus glia. (A) Relationship between cell type-specific SBS5 mutation rates and transcription levels (as estimated by single-cell RNA-seq [89]) in neurons (blue) and glia (orange). SBS5 mutation rates are shown relative to the value in the first bin for each cell type. 95% CIs were calculated by bootstrap resampling genes in each quartile 100 times. (B) Same as in (A), but for quartiles of genome accessibility, as calculated from genome-wide ATAC-seq read depths for the two cell types [90]. To obtain 95% CIs, we bootstrap resampled 500 kb windows in each accessibility quartile 100 times. (C) SBS5 mutation rates in ATAC-seq peaks [90] in neurons (blue) and glia (orange). Expected SBS5 mutation rates were obtained by randomly shuffling peak positions within the same chromosome and matching for GC% (within ± 1%) 100 times. 95% CIs were calculated by taking the 2.5 and 97.5 percentiles of the observed-to-expected SBS5 count ratio. Circles show results for SBS5 mutation rates and crosses for SBS5 + SBS40 mutation rates.

**Fig. S3:**
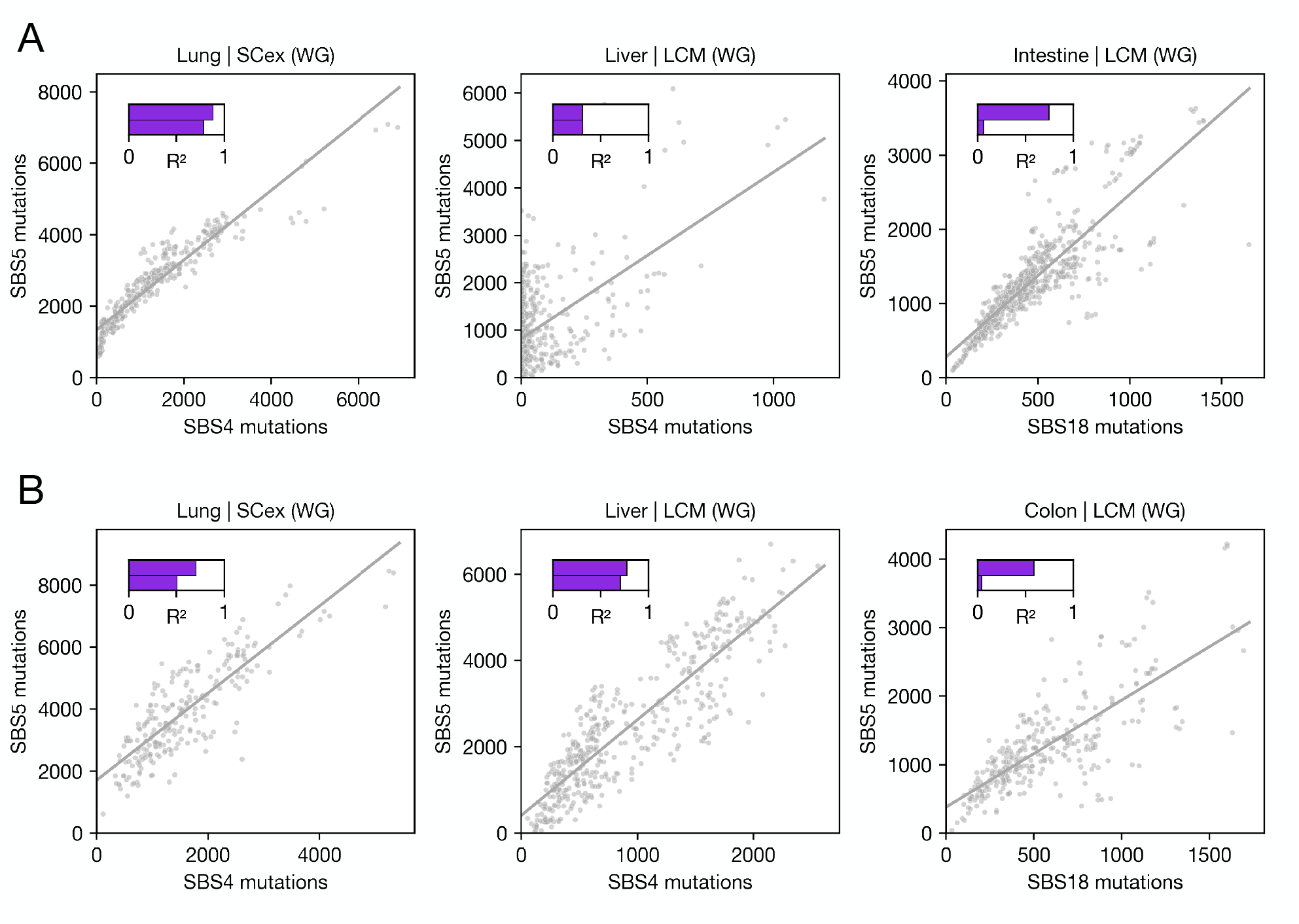
(A) Association between SBS5 and damage-specific signatures using the signature attributions in the original publications (data sources and signature attribution methods can be found in Table S3). The solid line represents the fit to all data points. The upper and lower bars show the semipartial *R*^2^ quantifying the variance explained by damage-specific signature in the models regressing SBS5 on the damage-specific signature and SBS5 on the damage-specific signature, SBS1, and age, respectively. (B) Same as in (A) but with signature attributions using SigProfiler (see Methods). Throughout all title panels: LCM: Laser Capture Microdissection, SCex: *in vitro* single-cell expansion, WG: whole-genome sequencing.

**Fig. S4:**
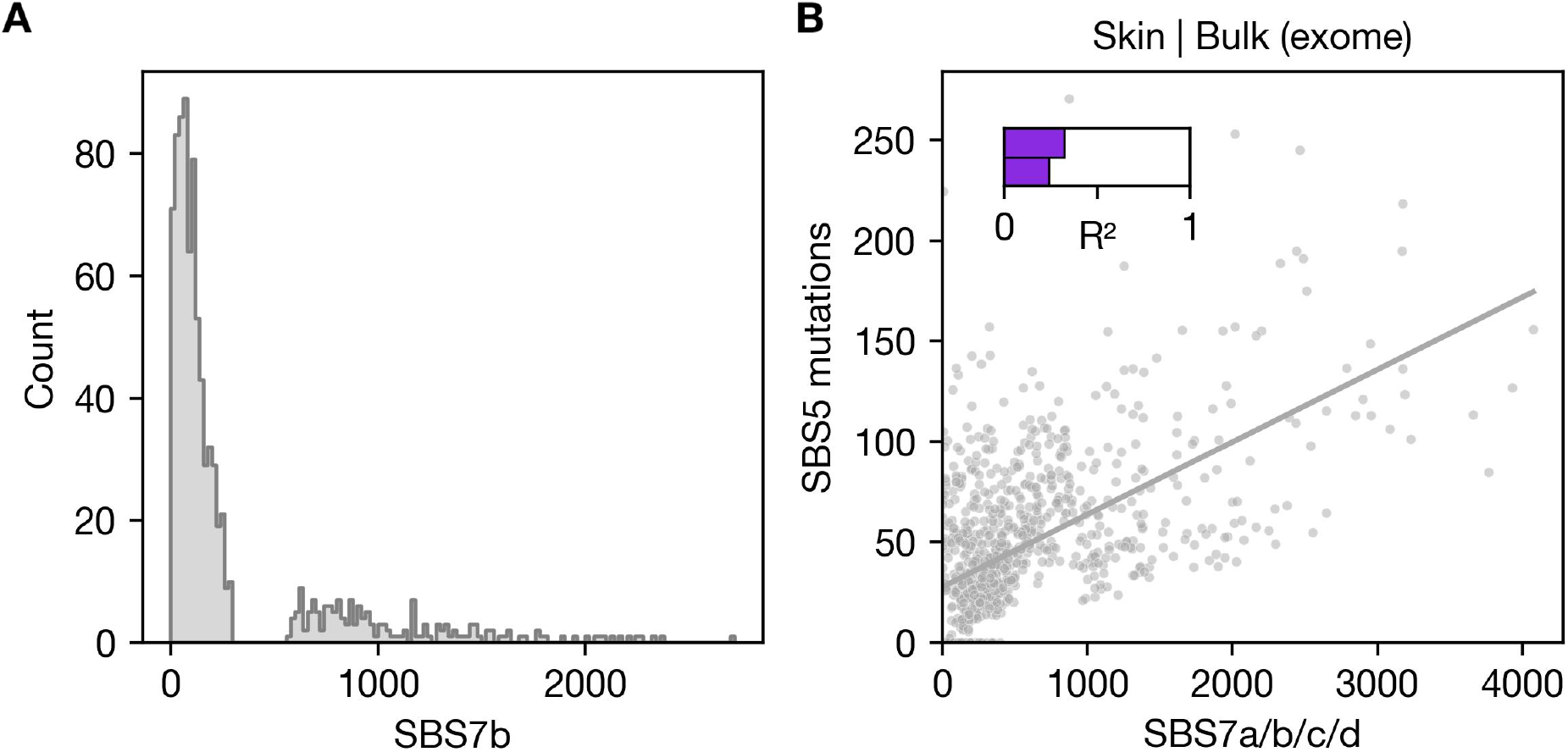
(A) Distribution of the number of SBS7 mutations attributed by SigNet [49] to skin microbiopsies in mutation data from [57]. (B) Association between SBS5 and the sum of SBS7a+b+c+d in skin microbiopsies. The solid line represents the fit to data from all cells. The upper and lower bars show the semipartial *R*^2^ quantifying the variance explained by SBS7a+b+c+d in the models regressing SBS5 on SBS7a+b+c+d and SBS5 on SBS7a+b+c+d, SBS1, and age, respectively.

**Fig. S5:**
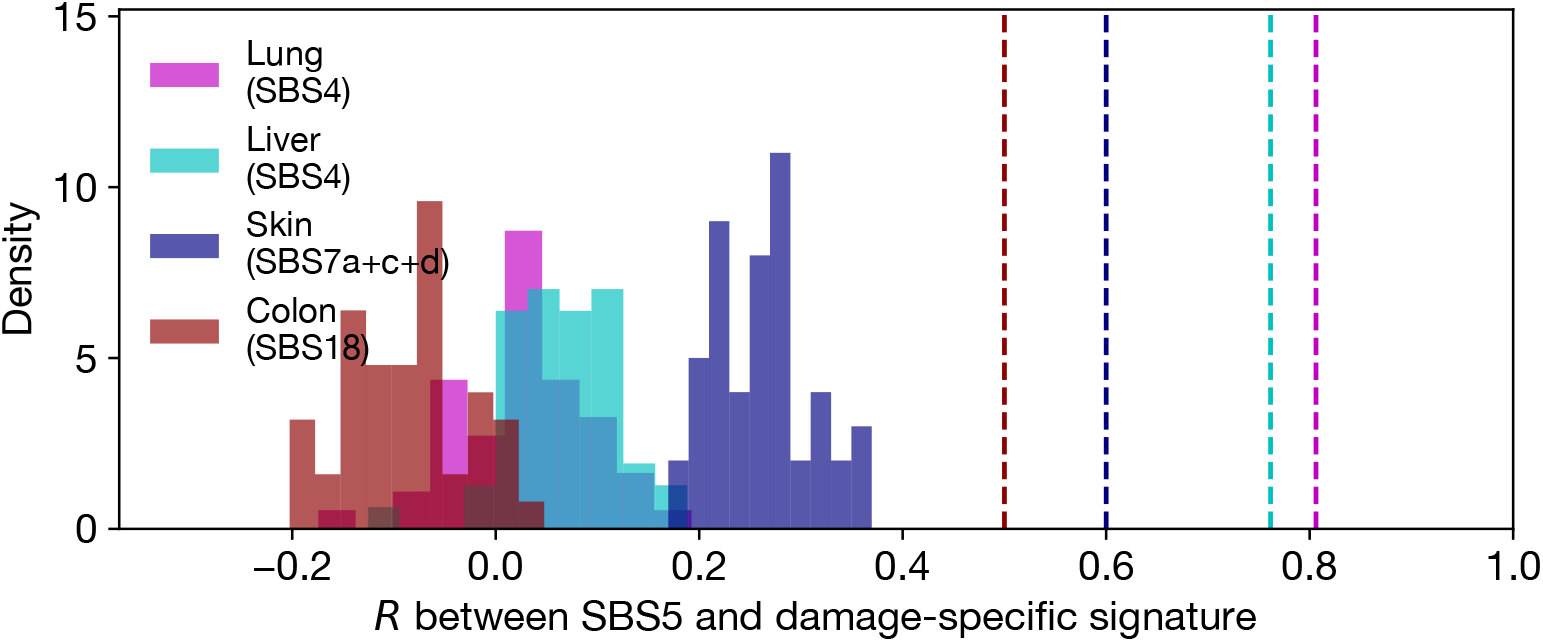
A test of whether signature attribution can introduce spurious correlations between signatures. We constructed synthetic datasets by shuffling the numbers of SBS5 and damage-specific signatures (as inferred from the data, and reported in Figure 3B-E), thus obtaining a comparable set of samples with no underlying correlation. We then inferred signature attribution in the synthetic data using SigNet. Reported here is the Pearson correlation coefficient between SBS5 and damage specific signatures. For reference, dashed lines denote the correlations found in real data (Figure 3B-E), which in all four cases are substantially and significantly greater than what is found in synthetic data.

**Fig. S6:**
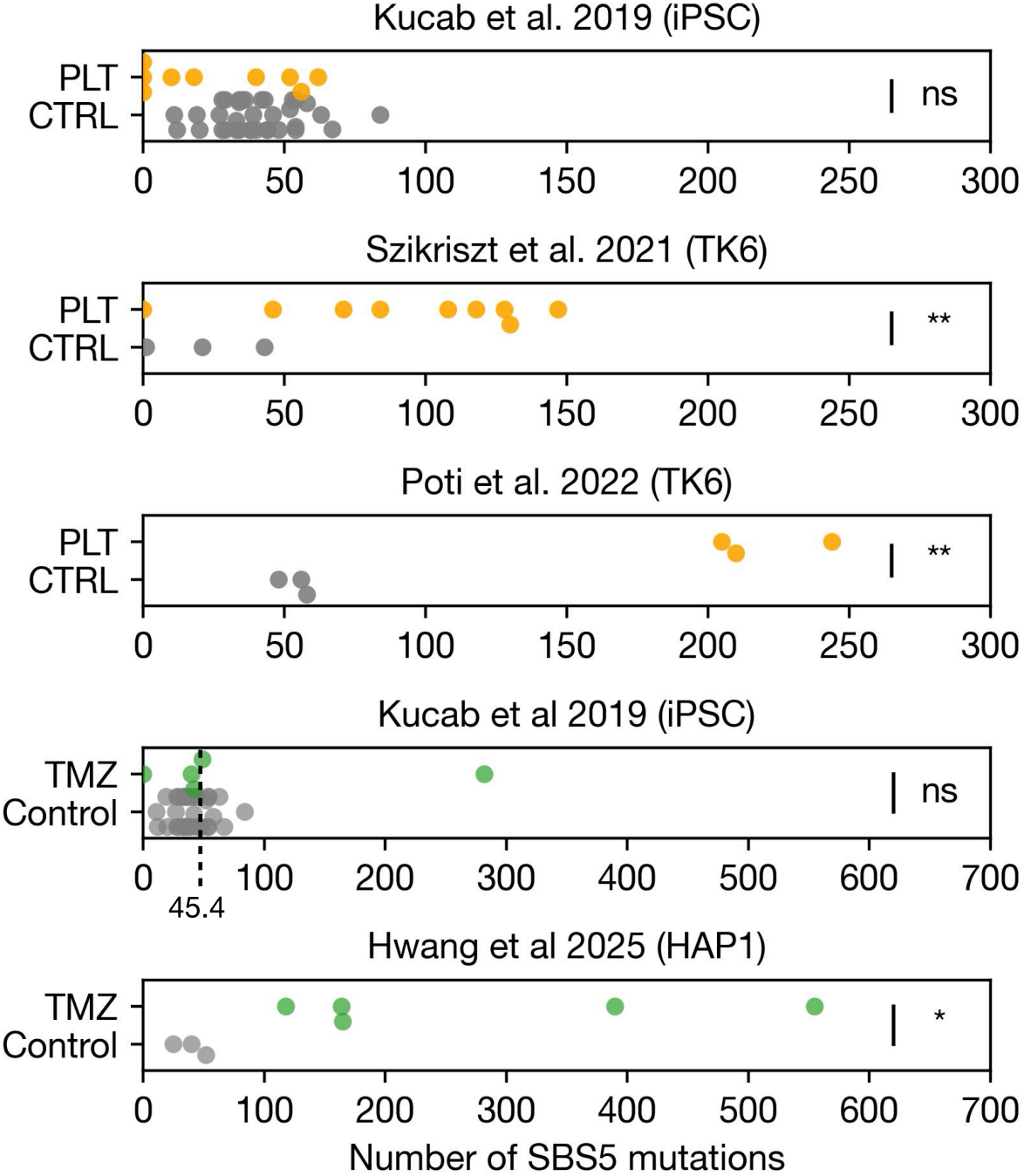
SBS5 mutation rates increase as a response to DNA damage in cell line experiments. Number of mutations attributed to SBS5 in cell lines (indicated in panel titles) following exposure to platinum-based chemotherapies (“PLT”, orange) or temozolomide (“TMZ”, green) compared to unexposed controls (“Control”, gray). Platinum-based agents (e.g., cisplatin, oxaliplatin) are grouped into a single category. To minimize differences with the original publications, mutation data was downloaded from the original sources and signatures attributions were inferred using SigProfilerAssignment [91]. Some of the mutations in Szikriszt et al [52] and Póti et al 2022 [43] are based on the same sequencing data, but were originally identified using different mutation calling pipelines. We note that the two cases that are not significant rely on iPSCs, pointing to a possible difference among cell lines. Significance levels (one-sided Welch’s t-test): ∗: *p <* 0.05, ∗∗: *p <* 0.005, ns: not significant.

**Fig. S7:**
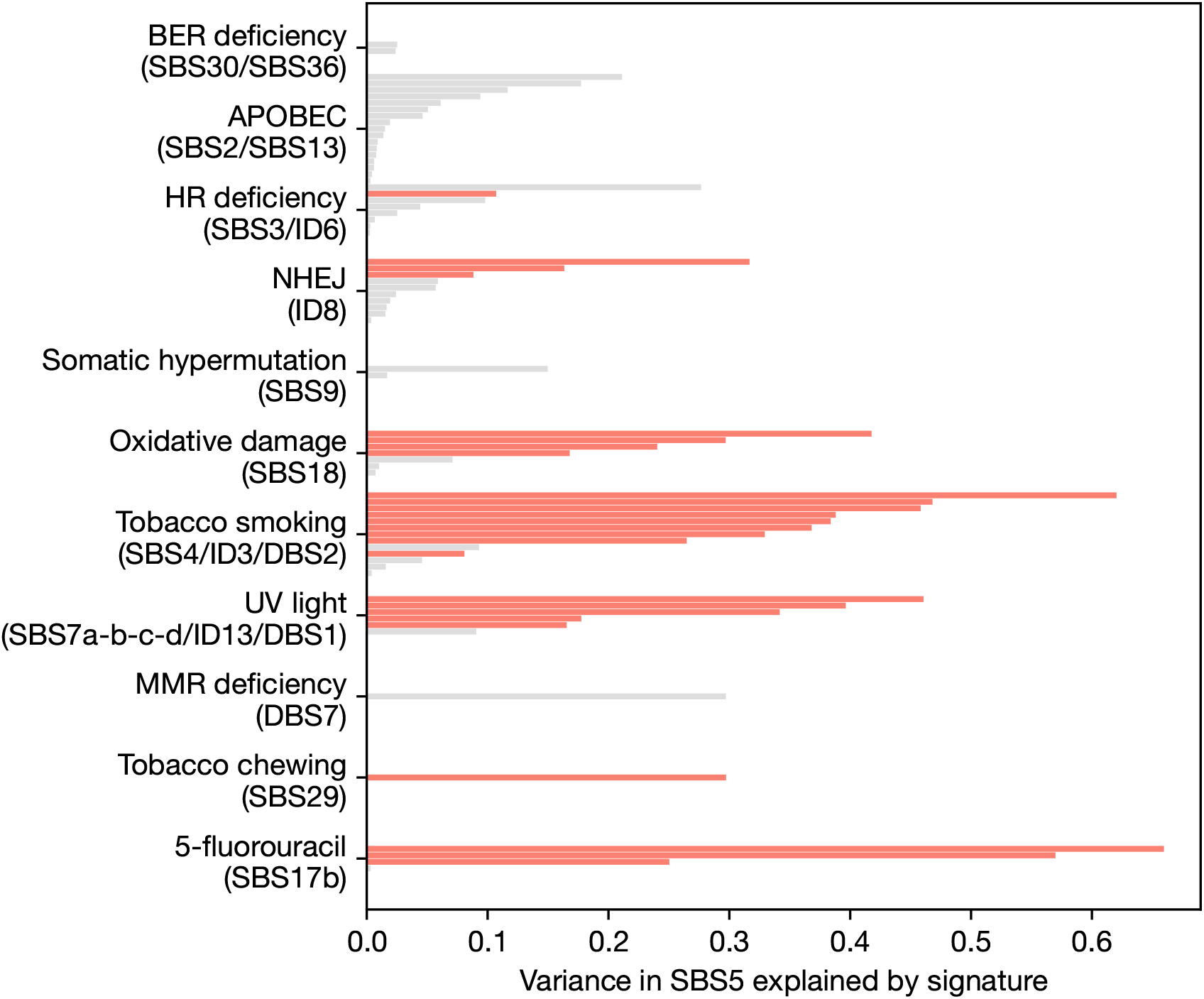
Association of SBS5 with other mutational signatures with a known etiology in PCAWG data. Semipartial R^2^ values quantify the variance in SBS5 activity attributable to individual mutational signatures (y-axis), after accounting for age, SBS1 (a proxy for cell division rate; see main text), tumor ploidy, and purity. Each bar represents a cancer type meeting the sample size threshold used in Figure 3 (see Methods). Signatures are ordered by the mean *R*^2^ value across studies. Red bars denote statistically significant associations with SBS5 (i.e., p <0.05 after Bonferroni correction across all tests).

**Fig. S8:**
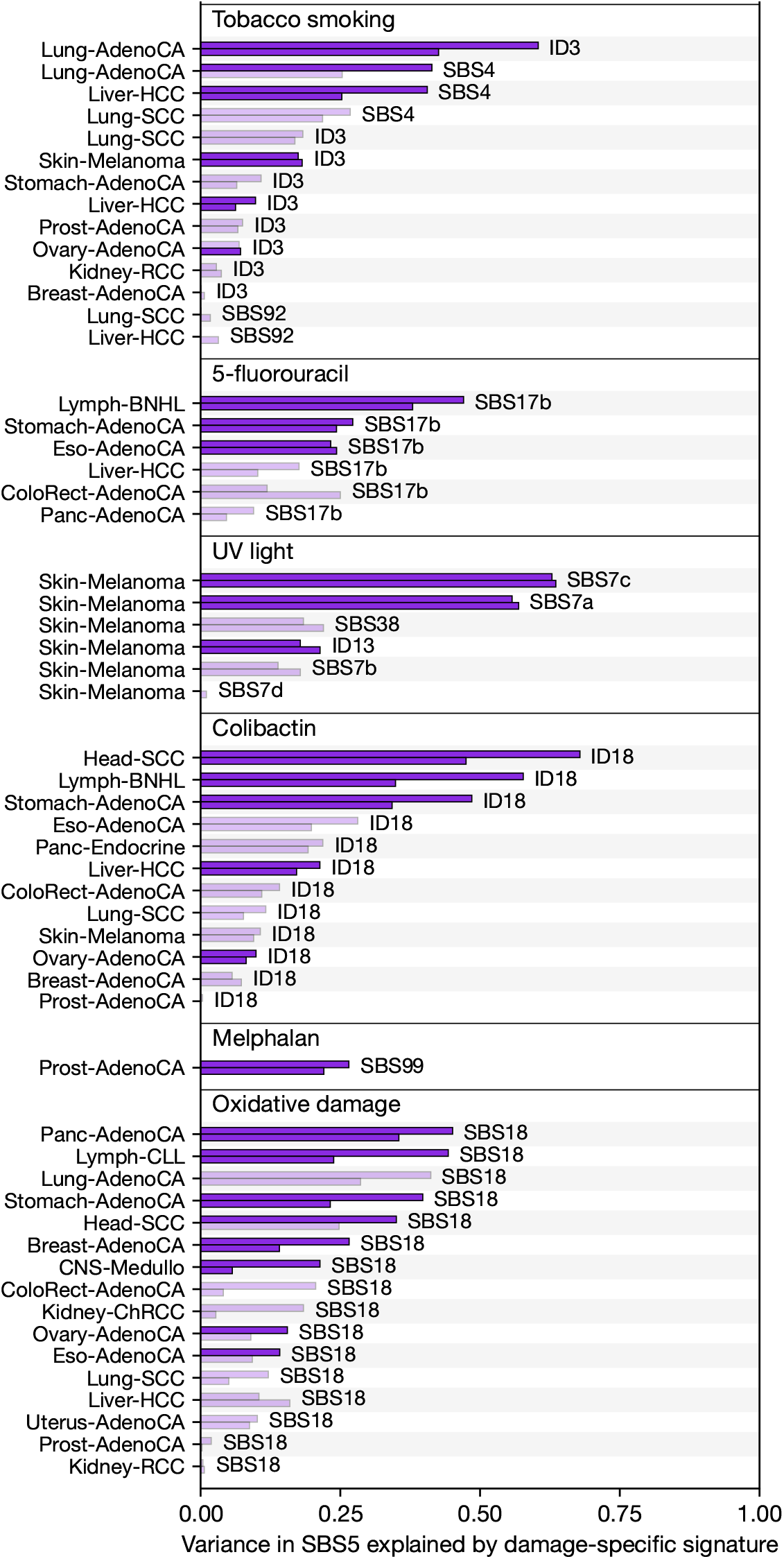
Association between SBS5 and various damage-specific signatures in PCAWG using signature attributions by MuSiCal, as provided in [35]. Semipartial *R*^2^ values quantifying the variance in SBS5 attributed to damage-specific signatures across tumors, with cancer types (left), signatures (right), and etiologies (top) indicated. The upper purple bars show the variance in SBS5 explained by a damage-specific signature in a baseline model with two covariates (ploidy and purity); lower bars represent the signature’s contribution to variance after also accounting for age and SBS1 (see Methods). Solid bars denote comparisons in which the damage-specific association with SBS5 was significant (*p <* 0.05 after Bonferroni correction). See Table S1 for data across all tested associations.

**Fig. S9:**
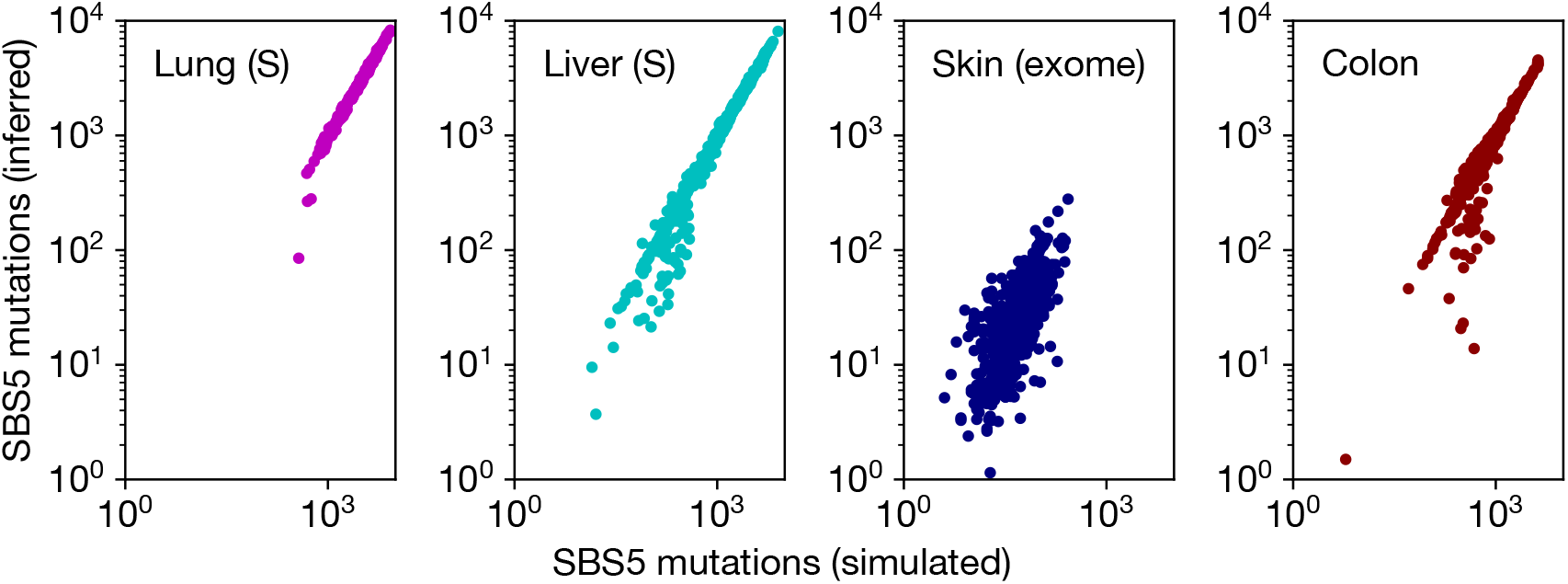
Test of the reliability of SBS5 inference across datasets used in Figure 3B-E. We constructed synthetic datasets by shuffling the numbers of SBS5 and damage-specific signatures (as inferred from the data, and reported in Figure 3B-E), thus obtaining a comparable set of samples with no underlying correlation. We then inferred signature attribution in the synthetic data using SigNet. We report the number of simulated and inferred SBS5 mutations.

**Fig. S10:**
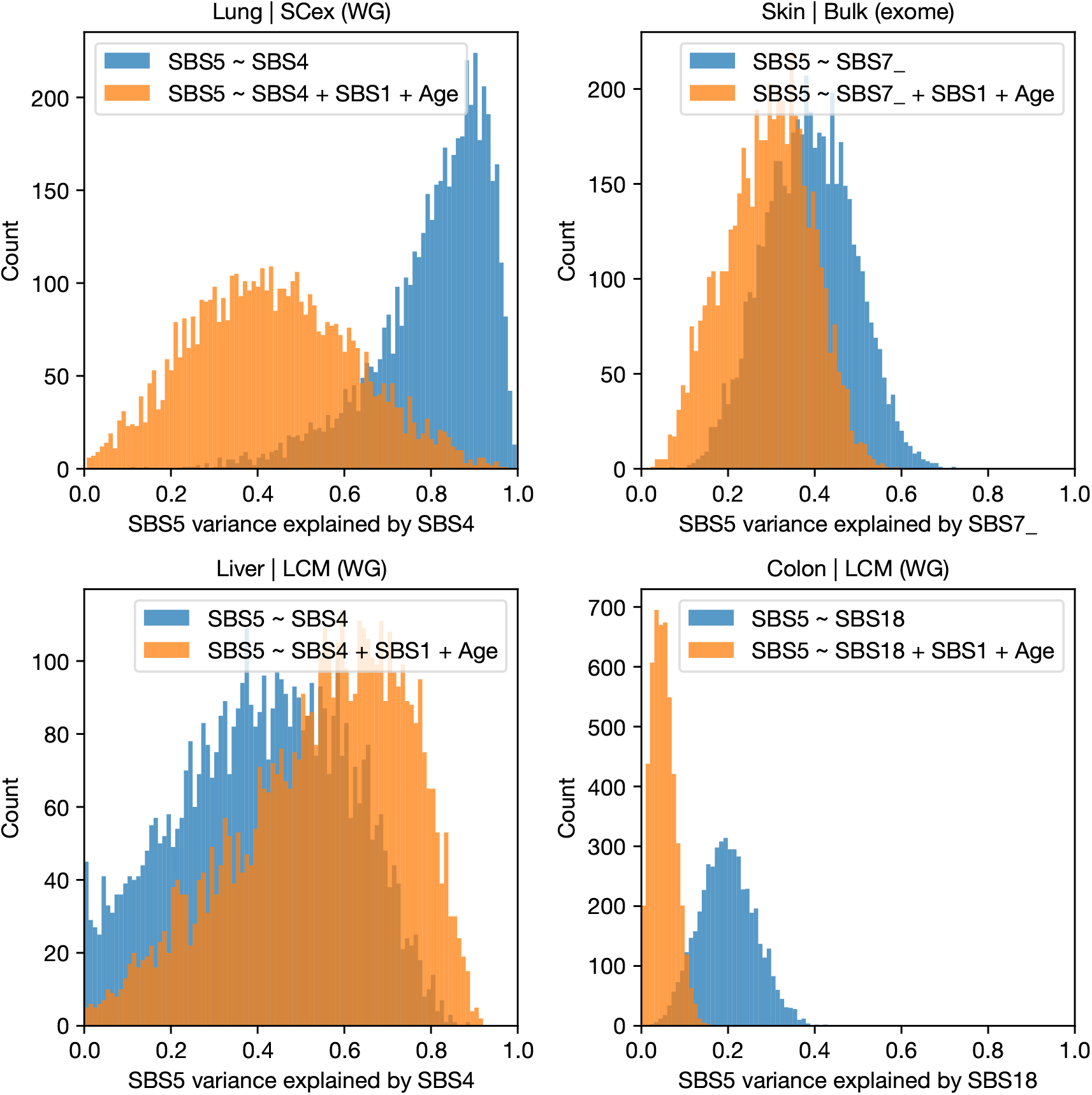
Distributions of semi-partial *R*^2^ quantifying the variance in SBS5 attributed to damage-specific signatures for 5,000 replicates, each of which resamples a single cell or microbiopsy per individual, for different OLS models (shown in the legend). Datasets are the same as in Figure 3B-E. Throughout all title panels: LCM: Laser Capture Microdissection, SCex: *in vitro* single-cell expansion, WG: whole-genome sequencing.

**Fig. S11:**
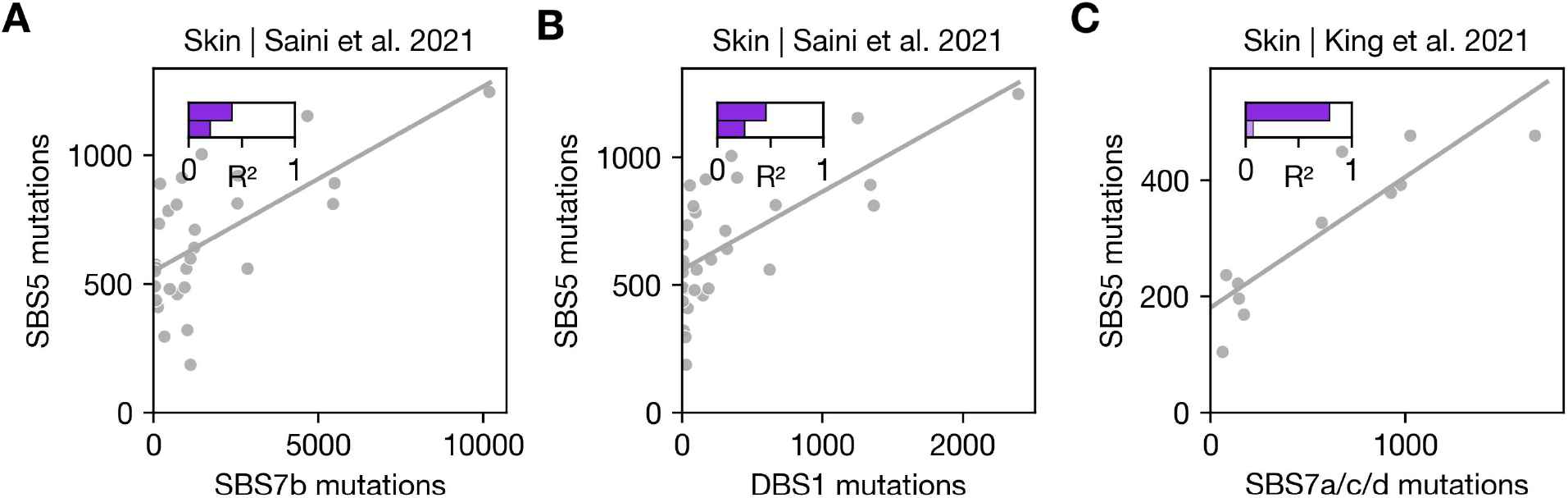
(A) Association between SBS5 and SBS7b in whole-genome sequences derived from colonies based on single skin cells, using the signature attributions in the original publication [58]. The solid line represents the fit to data from all cells. The upper and lower bars show the semipartial *R*^2^ quantifying the variance explained by SBS7b in the models regressing SBS5 on SBS7b and SBS5 on SBS7b, SBS1, and age, respectively. Solid bars denote comparisons in which the damage-specific association with SBS5 was significant (*p <* 0.05). (B) Same as in (A) but for mutations assigned to the damage-specific signature DBS. (C) Same as in (A) but for mutations assigned to SBS7a+c+d (i.e., the sum of all three signatures) in skin cell sequences from King et al. 2021.

**Fig. S12:**
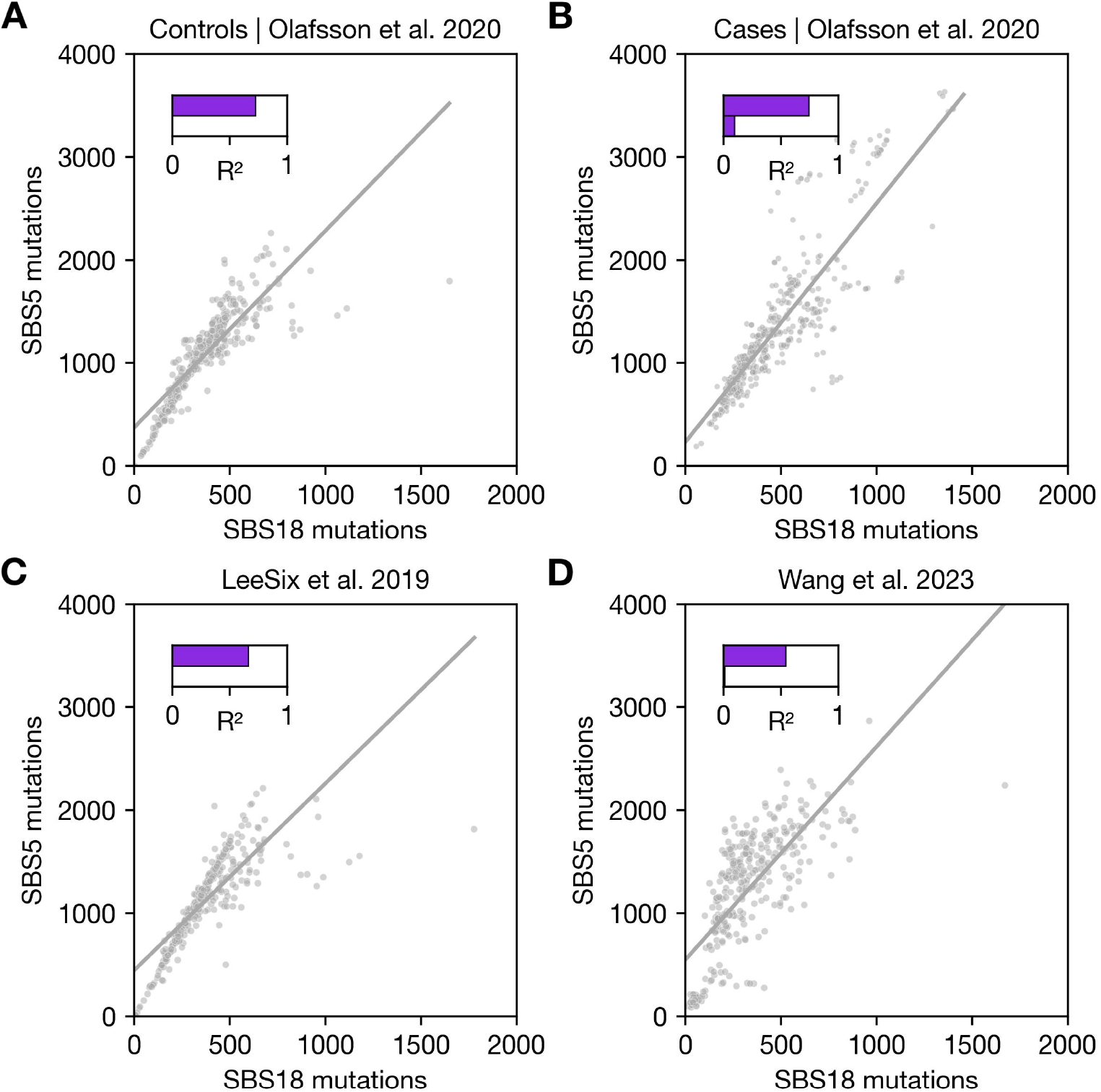
Associations between SBS18 and SBS5 using the signature attributions in the original publications (indicated in the panel titles). The solid lines represent the fit to all data points. The upper and lower bars show the semipartial R2 quantifying the variance explained by SBS18 in the models regressing SBS5 on SBS18 and SBS5 on SBS18, SBS1, and age, respectively (see Methods). (A) Data from non-IBD samples in [56] (B) Colon data from IBD individuals in [56]. (C) Colon data from individuals in [92]. Note the underlying sequencing data is the same as in (A). (D) Small bowel data from [60].

**Fig. S13:**
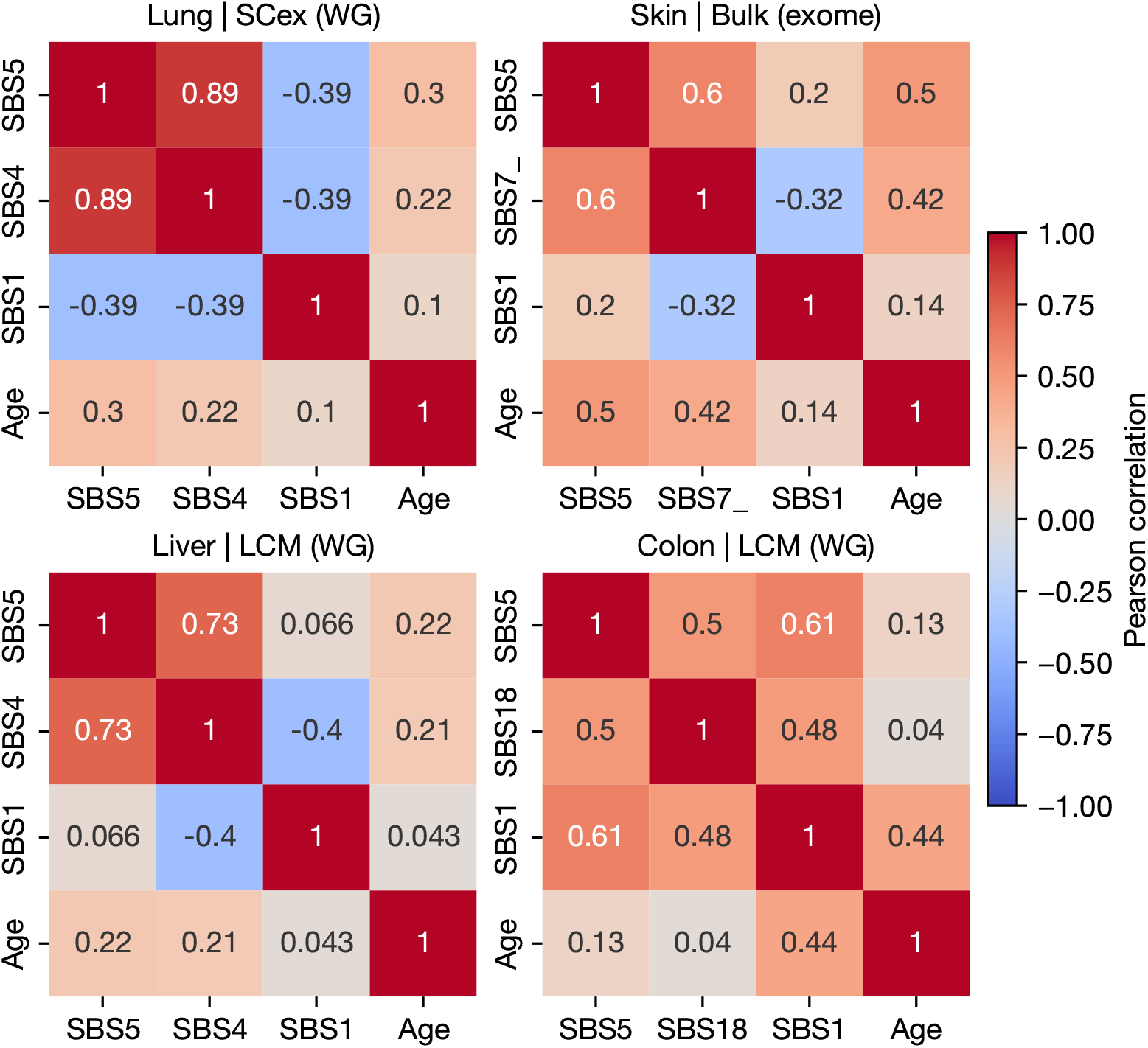
Pearson correlations between mutation counts assigned to damage-specific signatures, SBS1, SBS5, and age for the data displayed in Figure 3B-E. Throughout all title panels: LCM: Laser Capture Microdissection, SCex: *in vitro* single-cell expansion, WG: whole-genome sequencing.

**Fig. S14:**
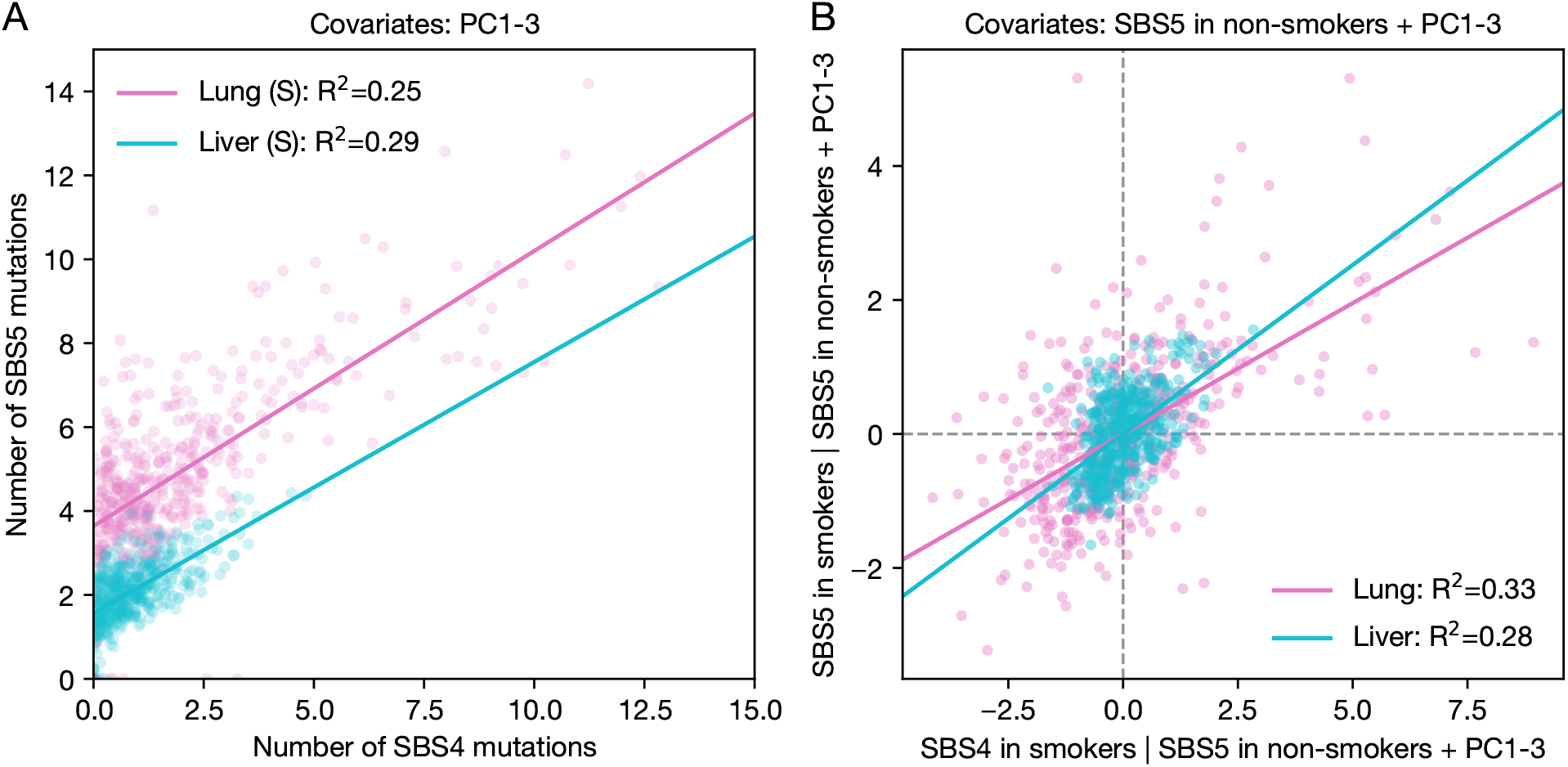
(A) Scatter plot of the association between the number of SBS4 and SBS5 mutations in 5 Mb genomic windows in lungs and livers of smokers. Signatures were attributed independently in each window, correcting for differences in mutational opportunities (see Methods). The solid line represents the linear regression fit to these data. In turn, the legend shows the proportion of variance explained by SBS4 rates *after accounting for the first three PCs* of a PC analysis of replication timing, GC content, gene density, and genome accessibility (see Methods). (B) Partial correlation between SBS4 and SBS5 across 5 Mb genomic windows in lung and liver of smokers, controlling for the first three PCs, as well as the SBS5 rate observed in non-smokers (considered as the baseline; see Methods). The solid line represents the linear regression fit, with the proportion of variance explained shown in the legend.

**Fig. S15:**
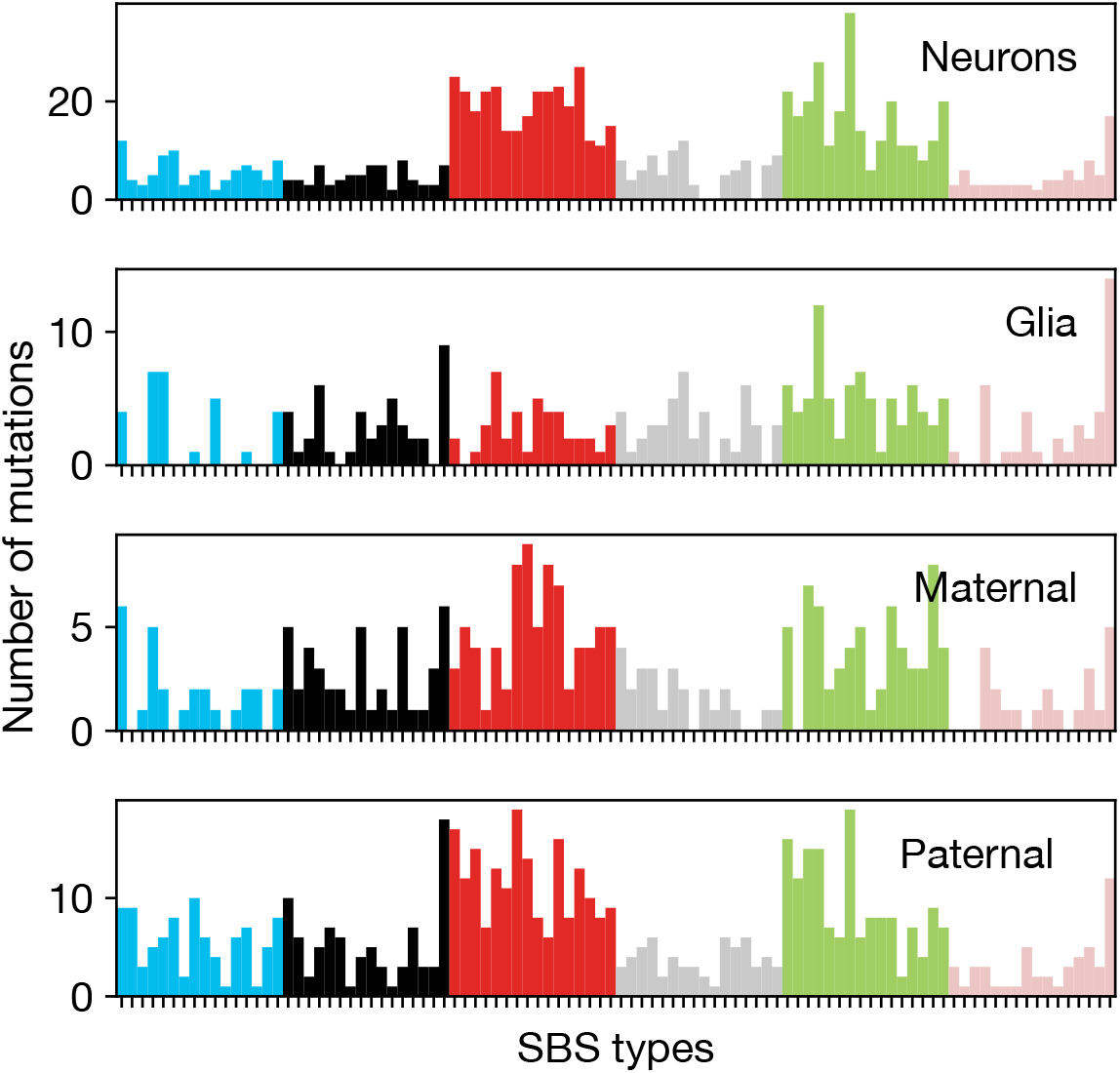
Distribution over 96 SBS types for clustered mutations in neurons, glia, maternal, and paternal mutations.

**Fig. S16:**
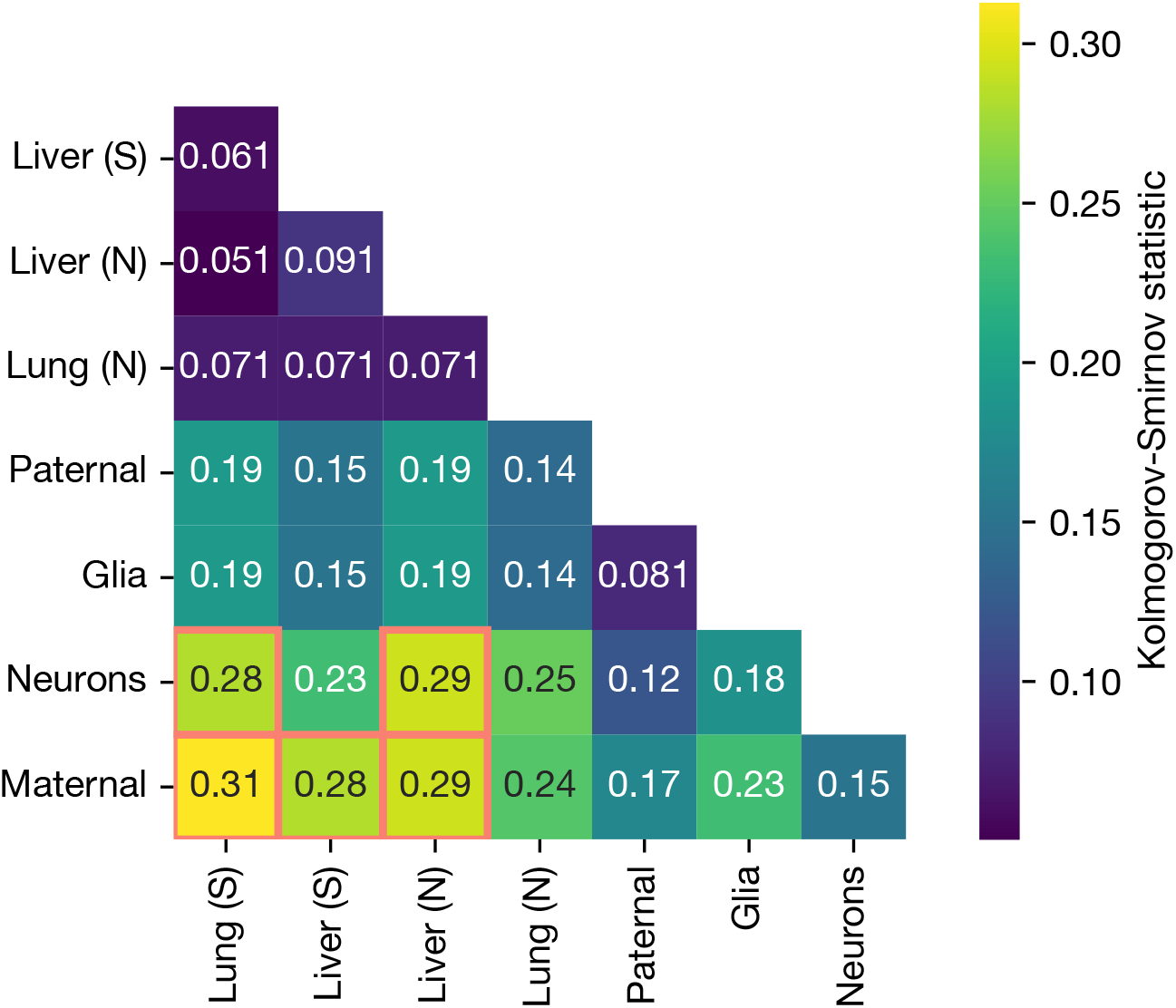
Kolmogorov-Smirnov statistics for all pairwise comparisons among the CCDFs in Figure 4B. Kolmogorov-Smirnov tests with a *p*-value below 0.05 (without correction for multiple testing) are indicated with red squares.

**Fig. S17:**
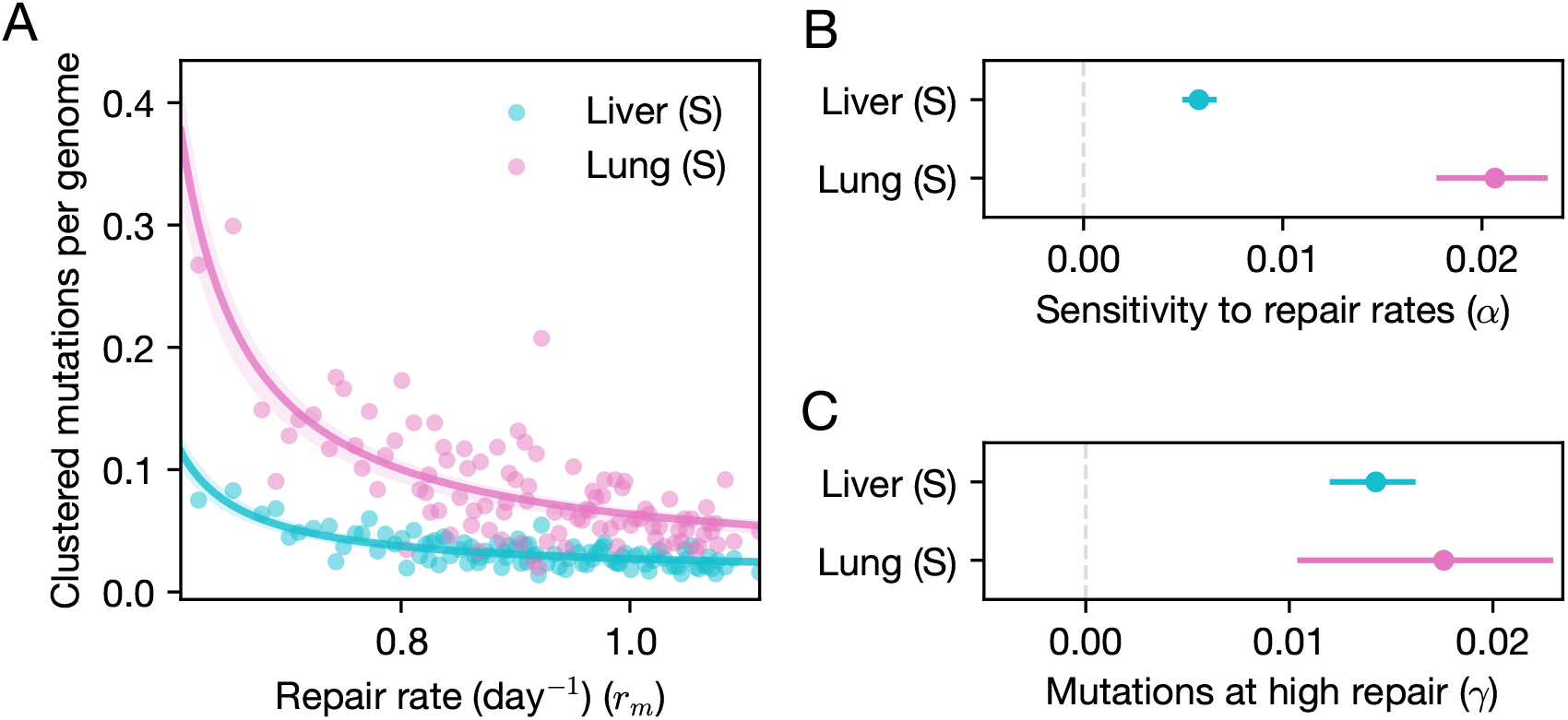
(A) Relationship between rates of NER (*x*-axis, see Methods) and the number of clustered mutations per genome (*y*-axis), for lungs and livers of smokers. Each point represents a percentile of the NER repair rate, estimated in 5 Mb windows along the genome. Lines represent the fit of the annotated model 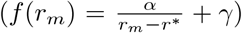. (**B**) Estimates of *α*. 95% CIs were obtained by bootstrapping individuals 1,000 times. (**C**) Same as in (B) but for *γ*. We note that the estimates of *α* and *γ* are not independent, as they were jointly fit to the data (see Methods).

**Fig. S18:**
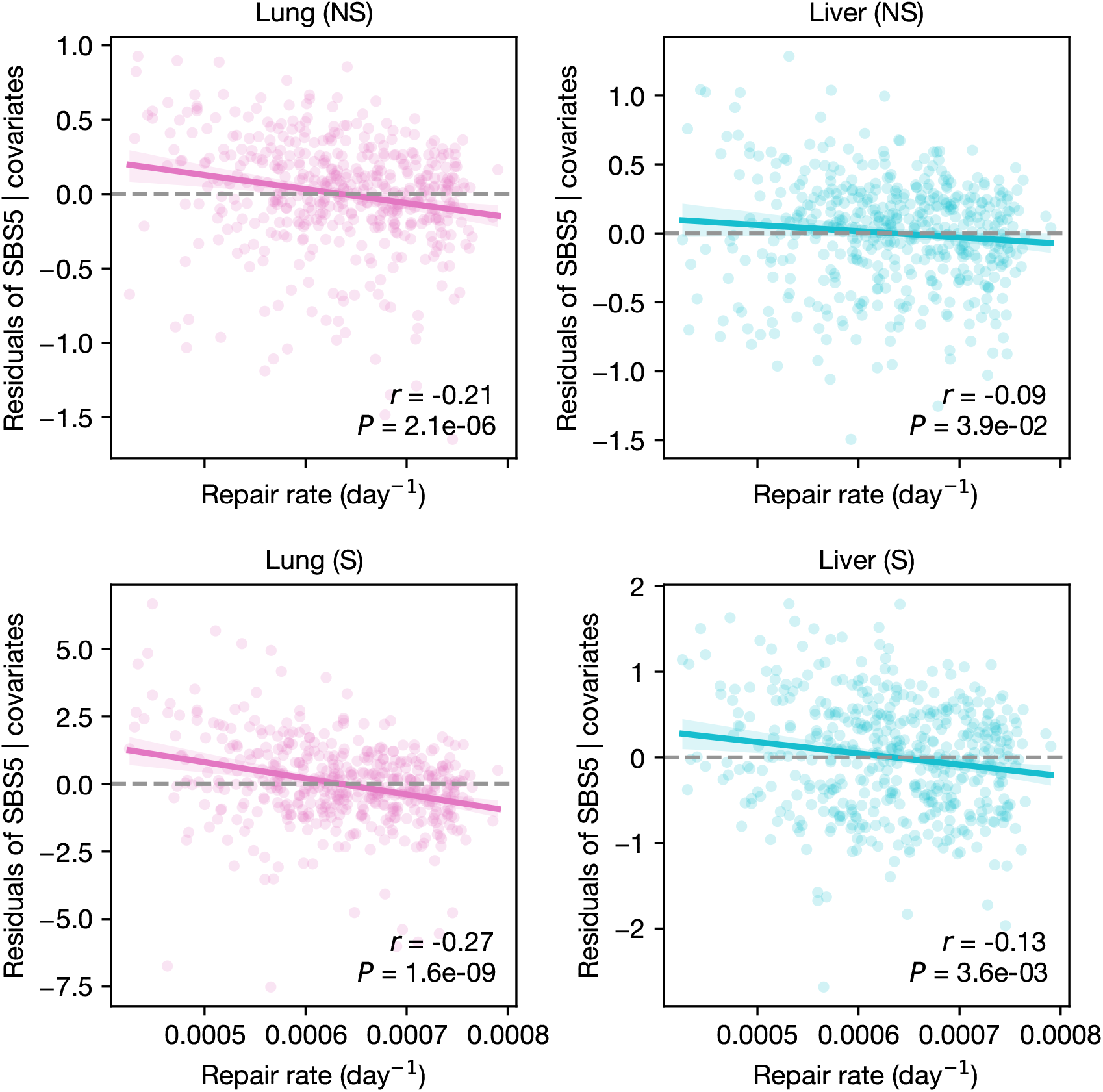
Associations between residual SBS5 mutation rates and NER repair rates across 5 Mb genomic windows. Signatures were attributed independently in each window, correcting for differences in mutational opportunities (see Methods). Residuals were obtained after accounting for the first three principal components (PCs) derived from a PC analysis of replication timing, genome accessibility, gene density, and GC content (see Methods). Pearson’s correlation coefficient and *p*-value are shown in the bottomright corner.

**Fig. S19:**
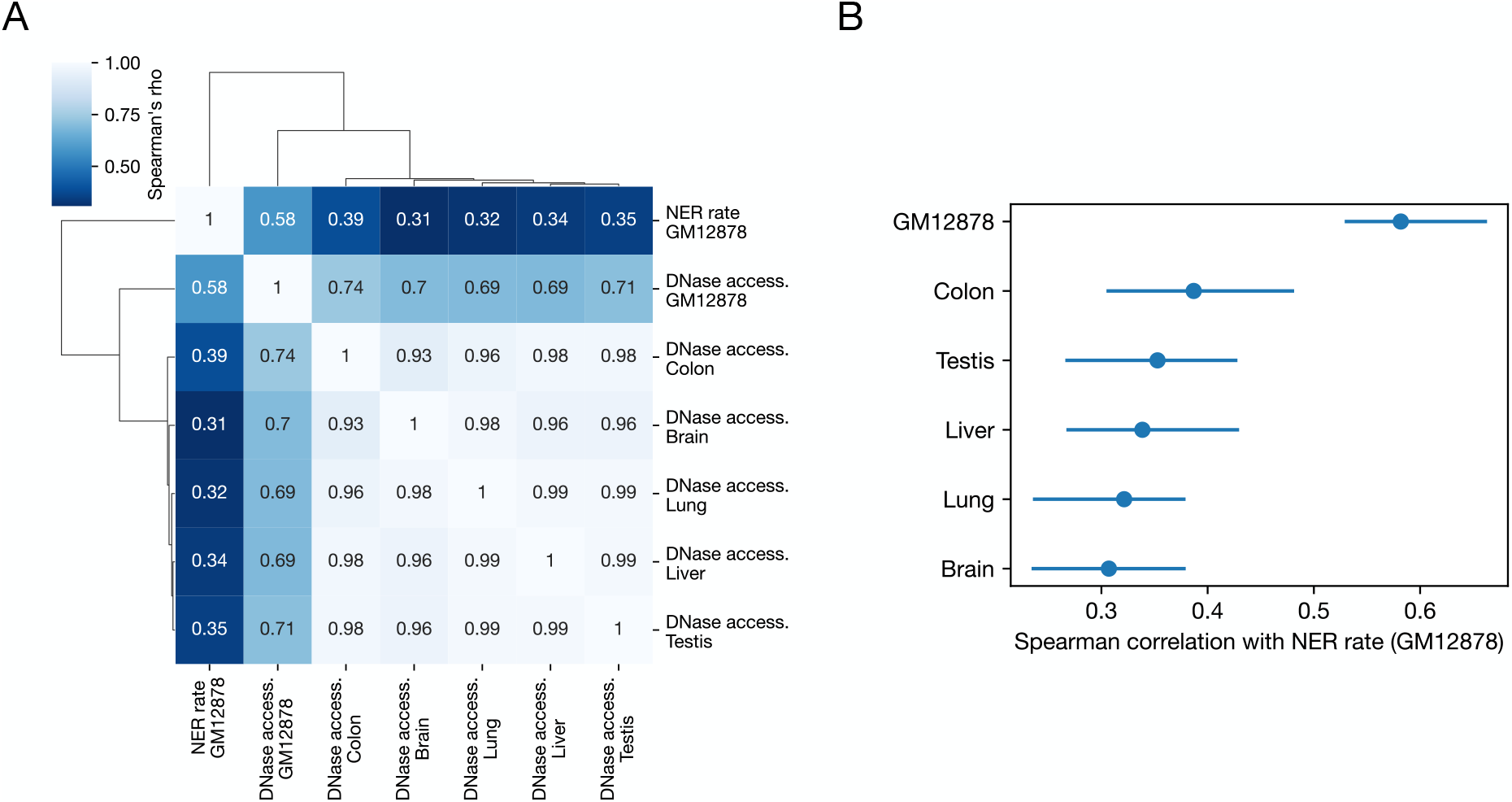
(A) Spearman correlations between NER rates (inferred from Damage-seq in GM12878 [66]; see Methods) and genome accessibilities (DNase-seq, ENCODE [93]) across 5 Mb windows. Rows and columns are ordered by hierarchical clustering (clustermap function in Python’s Seaborn). (B) Spearman correlations between NER rates and genome accessibility in each tissue. Horizontal lines represent the 95% CIs obtained by bootstrap resampling 5 Mb windows 500 times.

**Fig. S20:**
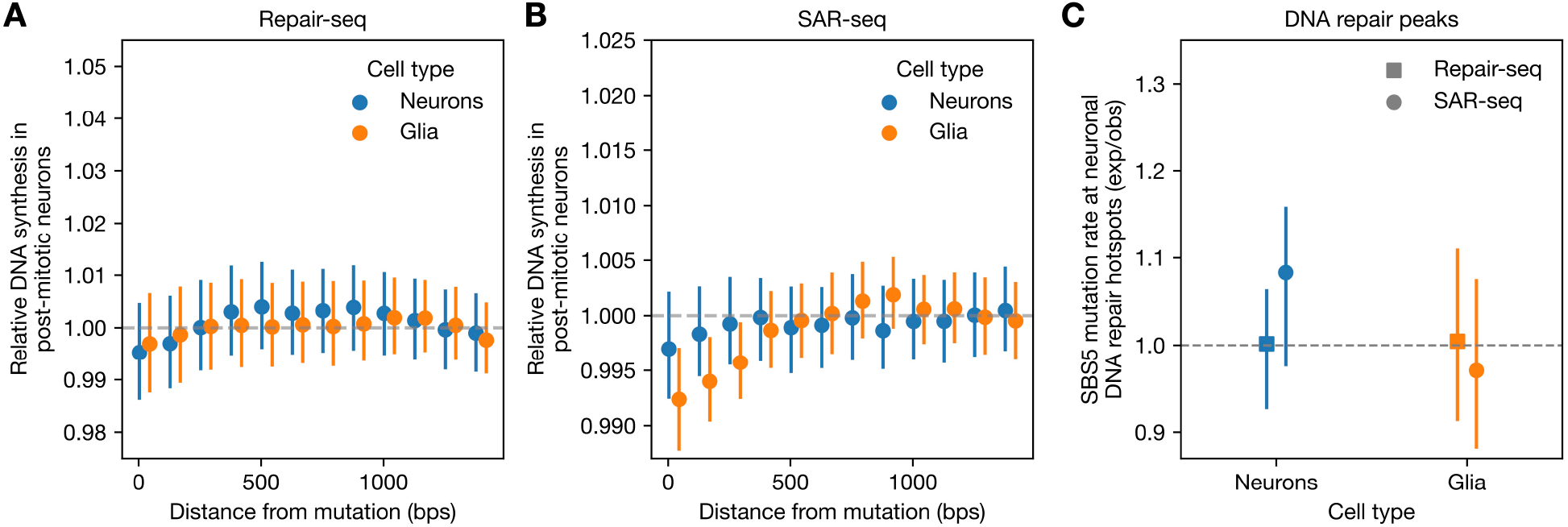
Local base composition effects cannot explain the observations in Figure 5F-H. (A) Depth of coverage of Repair-seq [63] as a function of the distance from mutation positions in neurons (blue) and glia (orange) shuffled and matched for their 5-mer context (see Methods). 95% CIs are computed by bootstrap resampling mutations 100 times. (B) Same as in (A) but for reads from SAR-seq [64]. (C) SBS5 mutation rates in neuronal DNA repair hotspots based on SAR-seq (circle) or Repair-seq (square) data, for mutations in neurons (blue) and glia (orange). Expected SBS5 mutation rates given the local base composition were obtained by randomly shuffling hotspot positions within the same chromosome and matching for GC% (within 1%) 100 times. Signature attributions were inferred by Signet. 95% CIs were calculated by taking the central 95%-tile of the observed-to-expected SBS5 count ratio.

**Fig. S21:**
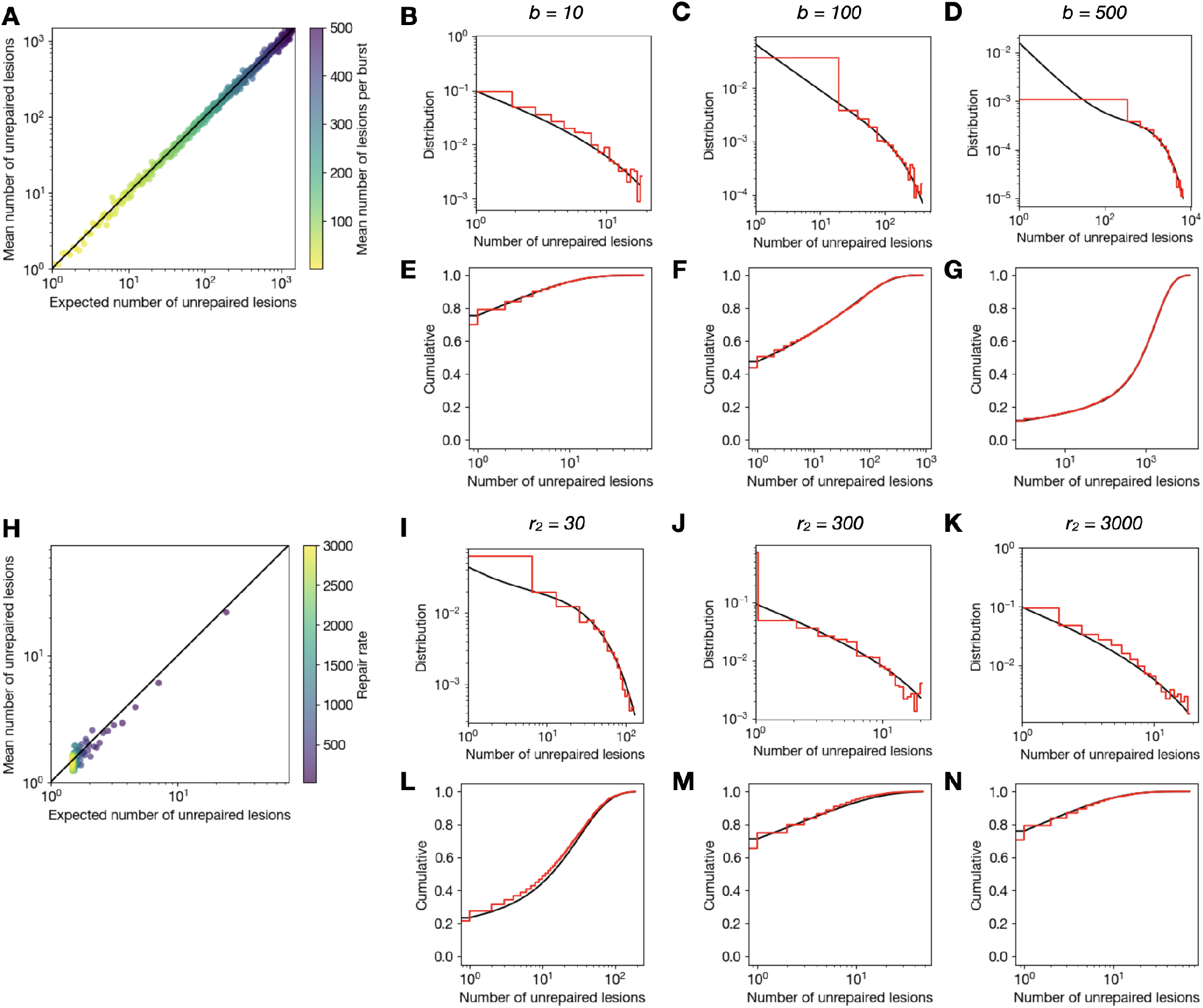
Comparison of model predictions with stochastic simulations. We present the results of varying the mean number of lesions per burst, *b* (A–G), and the repair rate, *r*_2_ (H–N). Unless otherwise noted, we set the simulation parameters to *f* = 20 per day, *N* = 10, *b* = 10, 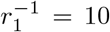 days, 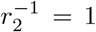 day. (A, H) Expected number of unrepaired lesions (equation 10) versus mean number of unrepaired lesions observed in the simulations across different parameter values. (B–D, I–K) Probability distribution of the number of unrepaired lesions. Analytical expectation (equation 9) in black, simulation results in red. (E–G, L–N) Cumulative distribution function of the number of unrepaired lesions. Analytical expectation (derived from equation 9) in black, simulation results in red.

**Fig. S22:**
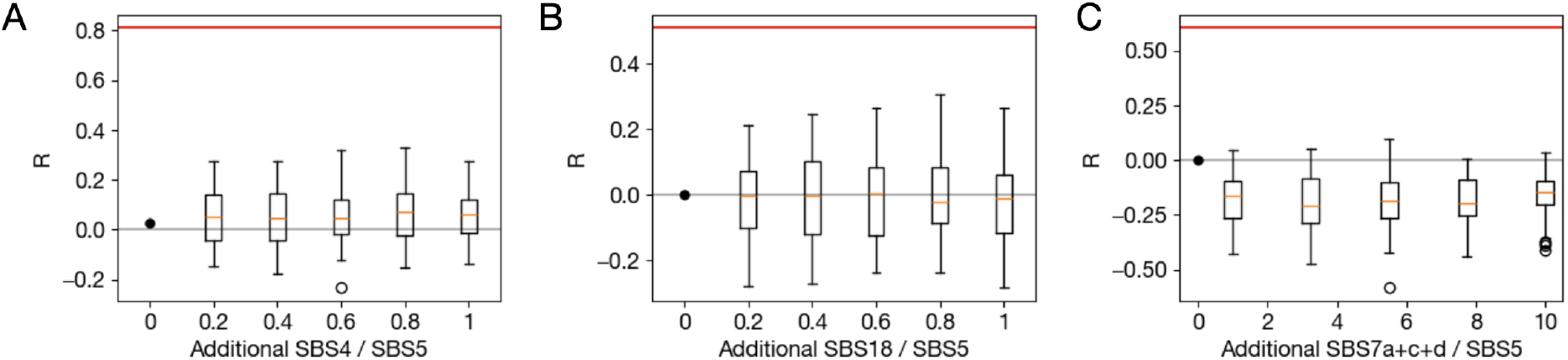
Pearson correlation coefficient between SBS5 and damage specific signatures as inferred from synthetic data with varying rates of damage specific signatures. The number of damage-specific mutations is expressed as a proportion of the mean number of SBS5 mutations, kept constant. Red lines correspond to Pearson correlation found in data (Figure 3B-E). (A) Additional SBS4 mutations are added to mutations found in lungs of non-smokers, consisting primarily of signature SBS5 (median of 1636 mutations per sample). (B) Additional SBS18 mutations were added to a background of SBS5 mutations (the number of SBS5 mutations was Poisson distributed with mean 1000). (C) Additional SBS7a+c+d mutations (signatures SBS7a, SBS7c and SBS7d in 10:1:1 proportion, as observed in the data) were added to a background of SBS5 mutations (the number of SBS5 mutations was Poisson distributed with mean 50).

**Fig. S23:**
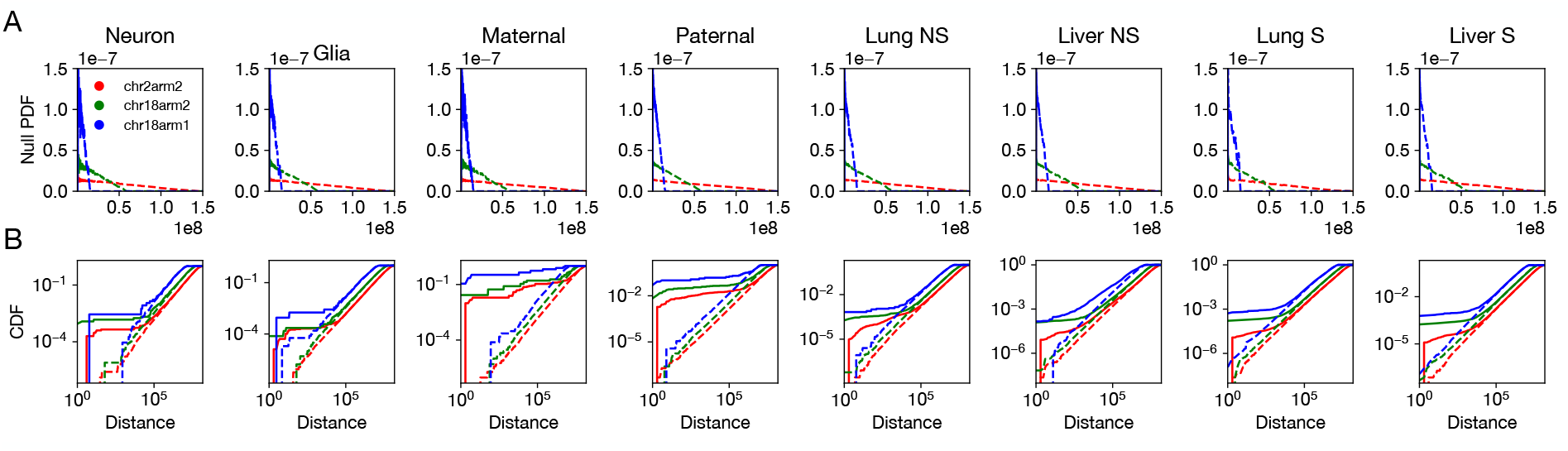
Distribution of distances for three chromosome arms (2*p*, 18*p*, 18*q* in red, green, blue) across datasets. (A) Probability density of distances between mutations computed between individuals. The distributions agree well with the null expectation (21), a triangular distribution determined by arm length (see Methods). (B) Cumulative distribution function (CDF) of distances between mutations computed within samples (solid lines) and between individuals (dashed line).

**Fig. S24:**
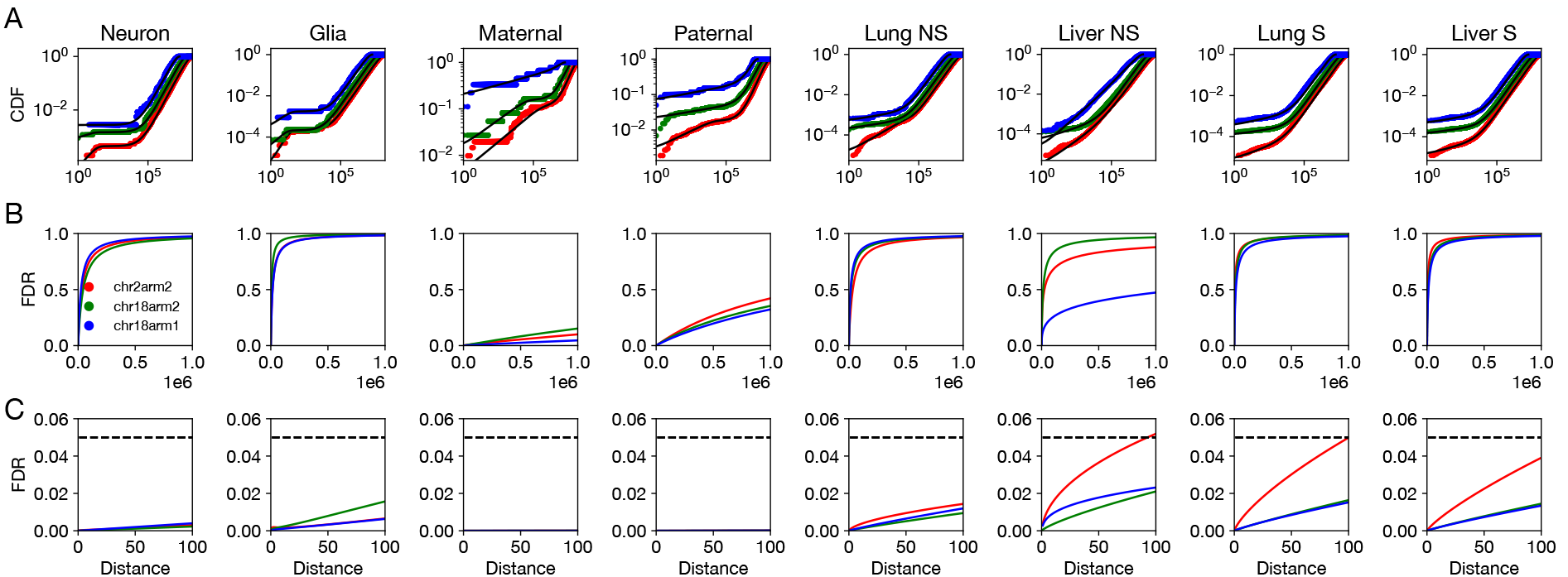
Modeling the distribution of distances for three chromosome arms (2*p*, 18*p*, 18*q* in red, green, blue) across datasets. (A) The cumulative distribution function (CDF) of distances between mutations computed within samples. Black lines show the best fit of the mixture distribution (22). (B) and (C) False discovery rate estimates for clustered mutations with thresholds at different distances, estimated from the fitted mixture distribution. We impose a threshold of 100 and exclude chromosome arms with FDR*>* 5% (dashed line in (C)).

**Fig. S25:**
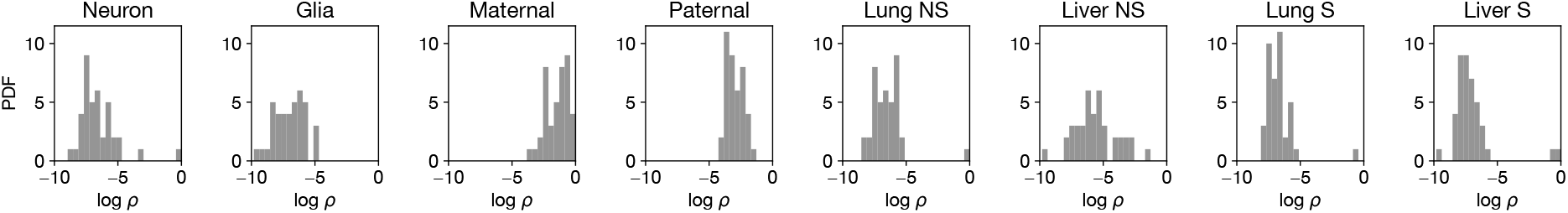
Distribution of cluster prevalences *ρ* (i.e. the fractions of pairs that are co-occurring, cf. eq. 22) inferred across chromosome arms for the datasets under study.

**Fig. S26:**
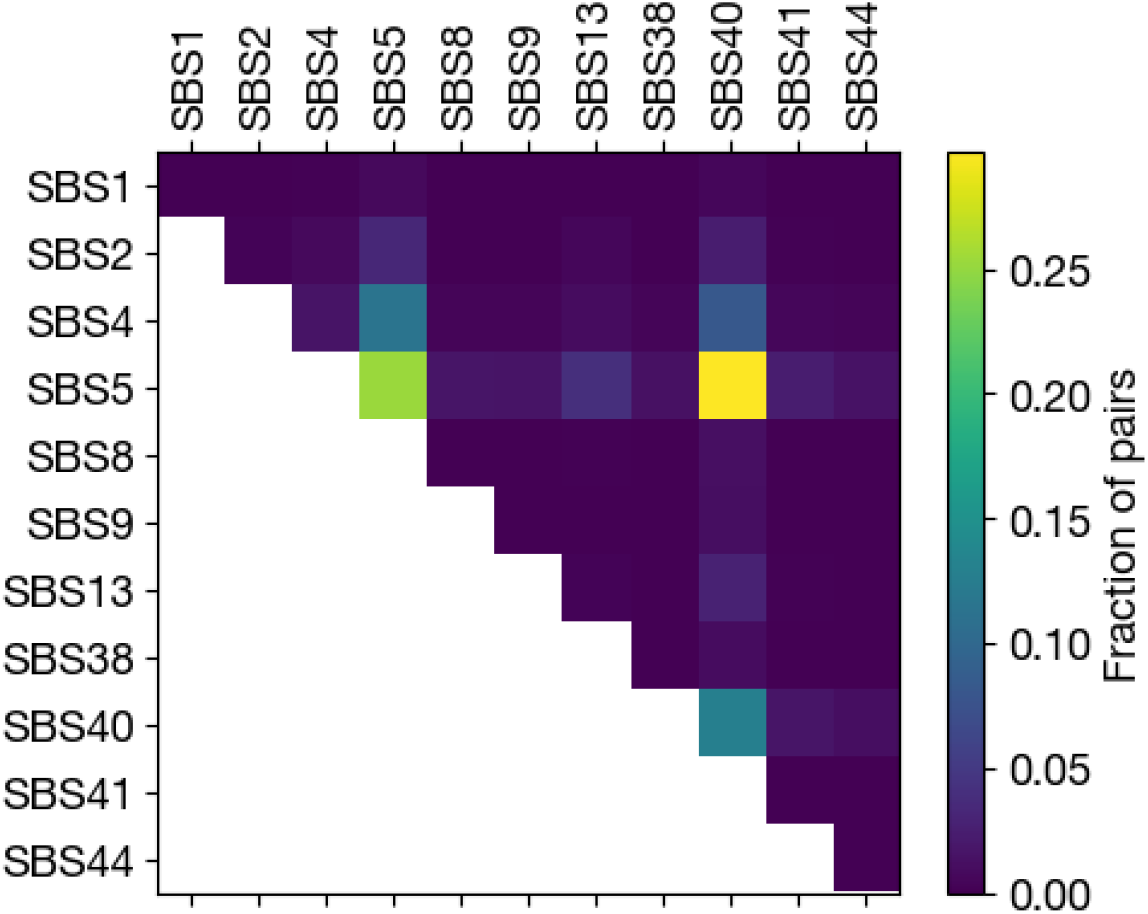
Estimated contribution of signatures to mutational clusters in lungs of smokers. Each cell in the heatmap represents the fraction of clustered mutation pairs attributed to the signature pair indicated by the corresponding row and column. For clarity, only the 11 signatures with the highest loadings are shown.

## References

[1] Alexandrov, L. B. et al. The repertoire of mutational signatures in human cancer. Nature 578, 94–101 (2020).

[2] Lodato, M. A. et al. Aging and neurodegeneration are associated with increased mutations in single human neurons. Science 359, 555–559 (2018).

[3] Spisak, N., de Manuel, M., Milligan, W., Sella, G. & Przeworski, M. The clock-like accumulation of germline and somatic mutations can arise from the interplay of DNA damage and repair. PLoS Biol. 22, e3002678 (2024).

[4] Pritchard, J. An owner’s guide to the human genome. https://web.stanford.edu/group/pritchardlab/HGbook.html (2023). Accessed: 2025-8-19.

[5] Stratton, M. R. Exploring the genomes of cancer cells: progress and promise. Science 331, 1553–1558 (2011).

[6] Alon, U. Systems medicine: Physiological circuits and the dynamics of disease (Chapman and Hall/CRC, Boca Raton, 2023).

[7] Koh, G., Degasperi, A., Zou, X., Momen, S. & Nik-Zainal, S. Mutational signatures: emerging concepts, caveats and clinical applications. Nat. Rev. Cancer 21, 619–637 (2021).

[8] Koh, G. C. C. et al. A redefined InDel taxonomy provides insights into mutational signatures. Nat. Genet. 1–10 (2025).

[9] Nik-Zainal, S. et al. Mutational processes molding the genomes of 21 breast cancers. Cell 149, 979–993 (2012).

[10] Alexandrov, L. B. et al. Signatures of mutational processes in human cancer. Nature 500, 415–421 (2013).

[11] Tate, J. G. et al. COSMIC: the catalogue of somatic mutations in cancer. Nucleic Acids Res. 47, D941–D947 (2019).

[12] Alexandrov, L. B. et al. Mutational signatures associated with tobacco smoking in human cancer. Science 354, 618–622 (2016).

[13] Jiang, Y. et al. Quantification and mapping of alkylation in the human genome reveal single nucleotide resolution precursors of mutational signatures. ACS Cent Sci 9, 362–372 (2023).

[14] Rahbari, R. et al. Timing, rates and spectra of human germline mutation. Nat. Genet. 48, 126–133 (2016).

[15] Moore, L. et al. The mutational landscape of human somatic and germline cells. Nature 597, 381–386 (2021).

[16] Alexandrov, L. B. et al. Clock-like mutational processes in human somatic cells. Nat. Genet. 47, 1402–1407 (2015).

[17] Abascal, F. et al. Somatic mutation landscapes at single-molecule resolution. Nature 593, 405–410 (2021).

[18] Cagan, A. et al. Somatic mutation rates scale with lifespan across mammals. Nature 604, 517–524 (2022).

[19] Beichman, A. C. et al. Evolution of the mutation spectrum across a mammalian phylogeny. Molecular Biology and Evolution 40, msad213 (2023).

[20] Waters, T. R. & Swann, P. F. Thymine-DNA glycosylase and G to a transition mutations at CpG sites. Mutat. Res. 462, 137–147 (2000).

[21] Gao, Z., Wyman, M. J., Sella, G. & Przeworski, M. Interpreting the dependence of mutation rates on age and time. PLoS Biol. 14, e1002355 (2016).

[22] Seplyarskiy, V. B. & Sunyaev, S. The origin of human mutation in light of genomic data. Nat. Rev. Genet. 22, 672–686 (2021).

[23] Tomkova, M. et al. Human DNA polymerase epsilon is a source of C to T mutations at CpG dinucleotides. Nat. Genet. 56, 2506–2516 (2024).

[24] Martínez-Jiménez, F. et al. Pan-cancer whole-genome comparison of primary and metastatic solid tumours. Nature 618, 333–341 (2023).

[25] Ganz, J. et al. Contrasting somatic mutation patterns in aging human neurons and oligodendrocytes. Cell (2024).

[26] Essuman, K. et al. Somatic mutation in human cerebellum illustrates neuron type-specific patterns of age-related mutation. bioRxiv 2026–02 (2026).

[27] COSMIC Human Cancer Signatures: Signature SBS1. URL https://cancer.sanger.ac.uk/signatures/sbs/sbs1/. Accessed: 2025-08.

[28] Zhu, T., Tong, H., Du, Z., Beck, S. & Teschendorff, A. E. An improved epigenetic counter to track mitotic age in normal and precancerous tissues. Nature Communications 15, 4211 (2024).

[29] Lal, A., Liu, K., Tibshirani, R., Sidow, A. & Ramazzotti, D. De novo mutational signature discovery in tumor genomes using SparseSignatures. PLoS Comput. Biol. 17, e1009119 (2021).

[30] Zou, X. et al. A systematic CRISPR screen defines mutational mechanisms underpinning signatures caused by replication errors and endogenous DNA damage. Nat Cancer 2, 643–657 (2021).

[31] COSMIC human cancer signatures: Signature SBS5. URL https://cancer.sanger.ac.uk/signatures/sbs/sbs5/. Accessed: 2025-08.

[32] Sasani, T. A. et al. Large, three-generation human families reveal post-zygotic mosaicism and variability in germline mutation accumulation. Elife 8 (2019).

[33] Halldorsson, B. V. et al. Characterizing mutagenic effects of recombination through a sequence-level genetic map. Science 363 (2019).

[34] Pancotti, C. et al. Unravelling the instability of mutational signatures extraction via archetypal analysis. Front. Genet. 13, 1049501 (2022).

[35] Jin, H. et al. Accurate and sensitive mutational signature analysis with MuSiCal. Nat. Genet. 56, 541–552 (2024).

[36] Senkin, S. et al. Geographic variation of mutagenic exposures in kidney cancer genomes. Nature 629, 910–918 (2024).

[37] Gronska-Peski, M., Srinivasa, A. & Evrony, G. D. Divergent somatic mutation patterns among human cerebellar neuron types. bioRxiv 2025–09 (2025).

[38] Chatterjee, N. & Walker, G. C. Mechanisms of DNA damage, repair, and mutagenesis. Environ. Mol. Mutagen. 58, 235–263 (2017).

[39] Friedberg, E. C., Walker, G. C., Siede, W. & Wood, R. D. DNA repair and mutagenesis (American Society for Microbiology Press, 2005).

[40] Shibutani, S., Takeshita, M. & Grollman, A. P. Insertion of specific bases during DNA synthesis past the oxidation-damaged base 8-oxodG. Nature 349, 431–434 (1991).

[41] Zou, X. et al. A systematic CRISPR screen defines mutational mechanisms underpinning signatures caused by replication errors and endogenous DNA damage. Nat Cancer 2, 643–657 (2021).

[42] Kucab, J. E. et al. A compendium of mutational signatures of environmental agents. Cell 177, 821–836.e16 (2019).

[43] Póti, Á., Szikriszt, B., Gervai, J. Z., Chen, D. & Szüts, D. Characterisation of the spectrum and genetic dependence of collateral mutations induced by translesion DNA synthesis. PLoS Genet. 18, e1010051 (2022).

[44] Gyüre, Z. et al. Spontaneous mutagenesis in human cells is controlled by REV1-polymerase zeta and PRIMPOL. Cell Rep. 42, 112887 (2023).

[45] Anderson, C. J. et al. Strand-resolved mutagenicity of DNA damage and repair. Nature 630, 744–751 (2024).

[46] Prakash, S. & Prakash, L. Translesion DNA synthesis in eukaryotes: a one-or two-polymerase affair. Genes Dev. 16, 1872–1883 (2002).

[47] Tubbs, A. & Nussenzweig, A. Endogenous DNA damage as a source of genomic instability in cancer. Cell 168, 644–656 (2017).

[48] Lindahl, T. & Barnes, D. Repair of endogenous DNA damage. Cold Spring Harbor symposia on quantitative biology 65, 127–134 (2000).

[49] Colome, C. S., Anton, O. C., Seplyarskiy, V. & Weghorn, D. Mutational signature decomposition with deep neural networks reveals origins of clock-like processes and hypoxia dependencies. bioRxiv 2023.12.06.570467 (2023).

[50] Yoshida, K. et al. Tobacco smoking and somatic mutations in human bronchial epithelium. Nature 578, 266–272 (2020).

[51] Hwang, T. et al. Comprehensive whole-genome sequencing reveals origins of mutational signatures associated with aging, mismatch repair deficiency and temozolomide chemotherapy. Nucleic Acids Res. gkae1122 (2024).

[52] Szikriszt, B. et al. A comparative analysis of the mutagenicity of platinum-containing chemotherapeutic agents reveals direct and indirect mutagenic mechanisms. Mutagenesis 36, 75–86 (2021).

[53] ICGC/TCGA Pan-Cancer Analysis of Whole Genomes Consortium. Pan-cancer analysis of whole genomes. Nature 578, 82–93 (2020).

[54] Li, Y. R. et al. The interplay between dormant mutated cells and tumor promotion by chronic tissue damage in determining cancer risk. bioRxiv 2024.01.24.577147 (2024).

[55] Ng, S. W. K. et al. Convergent somatic mutations in metabolism genes in chronic liver disease. Nature 598, 473–478 (2021).

[56] Olafsson, S. et al. Somatic evolution in non-neoplastic IBD-affected colon. Cell 182, 672–684.e11 (2020).

[57] Olafsson, S. et al. Effects of psoriasis and psoralen exposure on the somatic mutation landscape of the skin. Nat. Genet. (2023).

[58] Saini, N. et al. UV-exposure, endogenous DNA damage, and DNA replication errors shape the spectra of genome changes in human skin. PLoS Genet. 17, e1009302 (2021).

[59] King, C. et al. Somatic mutations in facial skin from countries of contrasting skin cancer risk. Nat. Genet. 55, 1440–1447 (2023).

[60] Wang, Y. et al. APOBEC mutagenesis is a common process in normal human small intestine. Nat. Genet. 55, 246–254 (2023).

[61] Supek, F. & Lehner, B. Clustered mutation signatures reveal that error-prone DNA repair targets mutations to active genes. Cell 170, 534–547.e23 (2017).

[62] Petljak, M. et al. Mechanisms of APOBEC3 mutagenesis in human cancer cells. Nature 607, 799–807 (2022).

[63] Reid, D. A. et al. Incorporation of a nucleoside analog maps genome repair sites in postmitotic human neurons. Science 372, 91–94 (2021).

[64] Wu, W. et al. Neuronal enhancers are hotspots for dna single-strand break repair. Nature 593, 440–444 (2021).

[65] Seeberg, E., Steinum, A. L., Nordenskjöld, M., Söderhäll, S. & Jernström, B. Strand-break formation in DNA modified by benzo[alpha]pyrene diolepoxide. quantitative cleavage by escherichia coli uvrABC endonuclease. Mutat. Res. 112, 139–145 (1983).

[66] Hu, J., Adebali, O., Adar, S. & Sancar, A. Dynamic maps of UV damage formation and repair for the human genome. Proc. Natl. Acad. Sci. U. S. A. 114, 6758–6763 (2017).

[67] García-Muse, T. & Aguilera, A. R loops: from physiological to pathological roles. Cell 179, 604–618 (2019).

[68] Saini, N. & Gordenin, D. A. Hypermutation in single-stranded dna. DNA repair 91, 102868 (2020).

[69] Thapar, U. & Demple, B. Deployment of dna polymerases beta and lambda in single-nucleotide and multinucleotide pathways of mammalian base excision dna repair. DNA repair 76, 11–19 (2019).

[70] Kochenova, O. V., Daee, D. L., Mertz, T. M. & Shcherbakova, P. V. Dna polymerase ζ-dependent lesion bypass in saccharomyces cerevisiae is accompanied by error-prone copying of long stretches of adjacent dna. PLoS genetics 11, e1005110 (2015).

[71] Martin, S. K. & Wood, R. D. Dna polymerase ζ in dna replication and repair. Nucleic acids research 47, 8348–8361 (2019).

[72] Kim, J. et al. Somatic ERCC2 mutations are associated with a distinct genomic signature in urothelial tumors. Nat. Genet. 48, 600–606 (2016).

[73] Barbour, J. A. et al. Ercc2 mutations alter the genomic distribution pattern of somatic mutations and are independently prognostic in bladder cancer. Cell Genomics 4 (2024).

[74] Ginno, P. A. et al. Single-mitosis dissection of acute and chronic DNA mutagenesis and repair. Nat. Genet. (2024).

[75] Riva, L. et al. The mutational signature profile of known and suspected human carcinogens in mice. Nat. Genet. 52, 1189–1197 (2020).

[76] Aitken, S. J. et al. Pervasive lesion segregation shapes cancer genome evolution. Nature 583, 265–270 (2020).

[77] Friedman, N., Cai, L. & Xie, X. S. Linking stochastic dynamics to population distribution: an analytical framework of gene expression. Phys. Rev. Lett. 97, 168302 (2006).

[78] Ochab-Marcinek, A. & Tabaka, M. Bimodal gene expression in noncooperative regulatory systems. Proceedings of the National Academy of Sciences 107, 22096–22101 (2010).

[79] Rodriguez, G. P. et al. Mismatch repair-dependent mutagenesis in nondividing cells. Proceedings of the National Academy of Sciences 109, 6153–6158 (2012).

[80] Gillespie, D.T. Exact stochastic simulation of coupled chemical reactions. The journal of physical chemistry 81, 2340–2361 (1977).

[81] Elowitz, M. & Bois, J. Biological circuit design. https://biocircuits.github.io/index.html (2021). Accessed: 2025-8-24.

[82] Jónsson, H. et al. Parental influence on human germline de novo mutations in 1,548 trios from iceland. Nature 549, 519–522 (2017).

[83] Díaz-Gay, M. et al. Assigning mutational signatures to individual samples and individual somatic mutations with sigprofilerassignment. Bioinformatics 39, btad756 (2023).

[84] Roberts, N. hdp: R package for the hierarchical dirichlet process. https://github.com/nicolaroberts/hdp.

[85] Blokzijl, F., Janssen, R., van Boxtel, R. & Cuppen, E. MutationalPatterns: comprehensive genomewide analysis of mutational processes. Genome Med. 10, 33 (2018).

[86] Kent, W. J. et al. The human genome browser at ucsc. Genome research 12, 996–1006 (2002).

[87] Pan-cancer analysis of whole genomes. Nature 578, 82–93 (2020).

[88] Li, H. Aligning sequence reads, clone sequences and assembly contigs with bwa-mem. arXiv preprint 1303.3997 (2013).

[89] Jeffries, A. M. et al. Single-cell transcriptomic and genomic changes in the ageing human brain. Nature 646, 657–666 (2025).

[90] Nott, A. et al. Brain cell type–specific enhancer–promoter interactome maps and disease-risk association. Science 366, 1134–1139 (2019).

[91] Diaz-Gay, M. et al. Assigning mutational signatures to individual samples and individual somatic mutations with SigProfilerAssignment. bioRxiv 2023.07.10.548264 (2023).

[92] Lee-Six, H. et al. The landscape of somatic mutation in normal colorectal epithelial cells. Nature 574, 532–537 (2019).

[93] Sloan, C. A. et al. Encode data at the encode portal. Nucleic acids research 44, D726–D732 (2016).

